# Modelling the niches of wild and domesticated Ungulate species using deep learning

**DOI:** 10.1101/744441

**Authors:** Mark Rademaker, Laurens Hogeweg, Rutger Vos

## Abstract

Knowledge of global biodiversity remains limited by geographic and taxonomic sampling biases. The scarcity of species data restricts our understanding of the underlying environmental factors shaping distributions, and the ability to draw comparisons among species. Species distribution models (SDMs) were developed in the early 2000s to address this issue. Although SDMs based on single layered Neural Networks have been experimented with in the past, these performed poorly. However, the past two decades have seen a strong increase in the use of Deep Learning (DL) approaches, such as Deep Neural Networks (DNNs). Despite the large improvement in predictive capacity DNNs provide over shallow networks, to our knowledge these have not yet been applied to SDM. The aim of this research was to provide a proof of concept of a DL-SDM^1^. We used a pre-existing dataset of the world’s ungulates and abiotic environmental predictors that had recently been used in MaxEnt SDM, to allow for a direct comparison of performance between both methods. Our DL-SDM consisted of a binary classification DNN containing 4 hidden layers and drop-out regularization between each layer. Performance of the DL-SDM was similar to MaxEnt for species with relatively large sample sizes and worse for species with relatively low sample sizes. Increasing the number of occurrences further improved DL-SDM performance for species that already had relatively high sample sizes. We then tried to further improve performance by altering the sampling procedure of negative instances and increasing the number of environmental predictors, including species interactions. This led to a large increase in model performance across the range of sample sizes in the species datasets. We conclude that DL-SDMs provide a suitable alternative to traditional SDMs such as MaxEnt and have the advantage of being both able to directly include species interactions, as well as being able to handle correlated input features. Further improvements to the model would include increasing its scalability by turning it into a multi-classification model, as well as developing a more user friendly DL-SDM Python package.

## 1 Introduction

### 1.1 Background

Biodiversity is in strong decline across the globe (*1, 2*). The main drivers are the loss and degradation of natural habitats through human activities (*3*). Loss of biodiversity negatively affects ecosystem functioning (*4*), and its conservation is therefore of high priority. However, knowledge of global biodiversity is still limited (*5*). This is partly due to the observation that the vast majority of known species occur in restricted ranges and low abundances (*6, 7*). Furthermore, data from areas with some of the highest biodiversity, such as the tropics, is relatively sparse (*8, 9*). Species distribution models (SDMs), which were initially developed in the early 2000s (*10, 11*), provide a partial solution to the scarcity of species data. SDMs relate patterns in the occurrence data to a selection of environmental predictors and use this information to predict the probability of presence outside of sampled areas (*12*). Predictions based on limited or geographically skewed input data, among other things, have implications for the quality and interpretation of model output (*13, 14*), and SDMs are subject to continuous improvement (*12*).

The MaxEnt software package (*15, 16*) is currently one of the most popular SDMs with *>* 1000 applications published since its introduction (*17*). The approach was originally developed to estimate the density of presences across the landscape (*15*). In the absence of knowledge on absolute population sizes, it provides a relative occurrence rate (ROR) per grid cell as output (*18*). However, for many species the available records cannot be seen as a random sample from the landscape, and the output will therefore not meet the assumptions for density estimation (*17*). Alternatively, using MaxEnt to predict the probability of presence in a cell requires a logistic transformation of the ROR (*16*). However, this transformation has also been criticized (*17, 19*). It includes a parameter τ, representing the background probability of presence for ‘average ‘presence localities. The value of τ has a large influence on the predicted output probabilities, but is arbitrarily set, rather than being fitted from the data (*20*). Considering the challenges in model interpretation when estimating density or probability of presence, MaxEnt is often used in a qualitative way by interpreting the output as an index of habitat suitability (*21, 22*).

In this research we propose an alternative approach for constructing SDMs, based on Deep Learning (DL). The past two decades have seen a strong increase in the use of Deep Learning (DL) (*23*), which has been attributed mainly to increased chip processing abilities, lower hardware costs and advances in machine learning algorithms (*24, 25*). DL is a subfield of machine learning that focuses on learning high-level abstractions in data (*25*). This is achieved by using a hierarchical architecture consisting of multiple interconnected layers, which in in turn contain multiple artificial neurons. A common type of DL is the application of Deep Neural Networks (DNNs). DNNs contain *>*2 layers and three basic computations are performed in each of them (Fig.1). (1) The neurons in a given layer receive input values from each of the neurons in the preceding layer. For the neurons in the first layer this means that they receive the raw values for each of the input variables. Each of these input values are multiplied by a specific weight, obtained through optimization, (2) the weighted input values are subsequently summed, and (3) the weighted sum is transformed using a non-linear activation function, which is selected from a set of candidate function by comparing the network’s performance using each of them. The transformed output value is passed on as input to the neurons in the next layer (*26*). Thus, what is learned by the network is the optimal set of weights for all the connections between neurons in adjoining layers, maximizing network performance. Each individual neuron is able to focus on a specific pattern in the data. For example, a neuron in the first layer might put most weight on all variables related to seasonality, and another neuron in the first layer assigns most weight to variables related to terrain and vegetation. A neuron in the second layer might then put most weight on the outputs of these particular two neurons in the first layer and thereby model the abstract concept of “seasonal lowland forest”. For classification purposes, the number of neurons in the final layer equals the number of classes to predict. The output of the neurons in the final layer are passed through a softmax function (*27*), which transforms them into probabilities that sum to 1 (eqn. 1). Although shallow networks containing a single hidden layer have been available in SDMs (*28, 29*), these typically ranked at the bottom in terms of performance (*30, 31*). Harris (*32*), used a two-layer network for SDM and already noticed a large increase in performance compared to single layered models. Current methods will allow us to create much deeper models still and further improve performance.

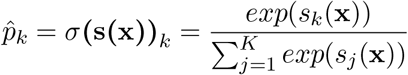

**Figure 1:**
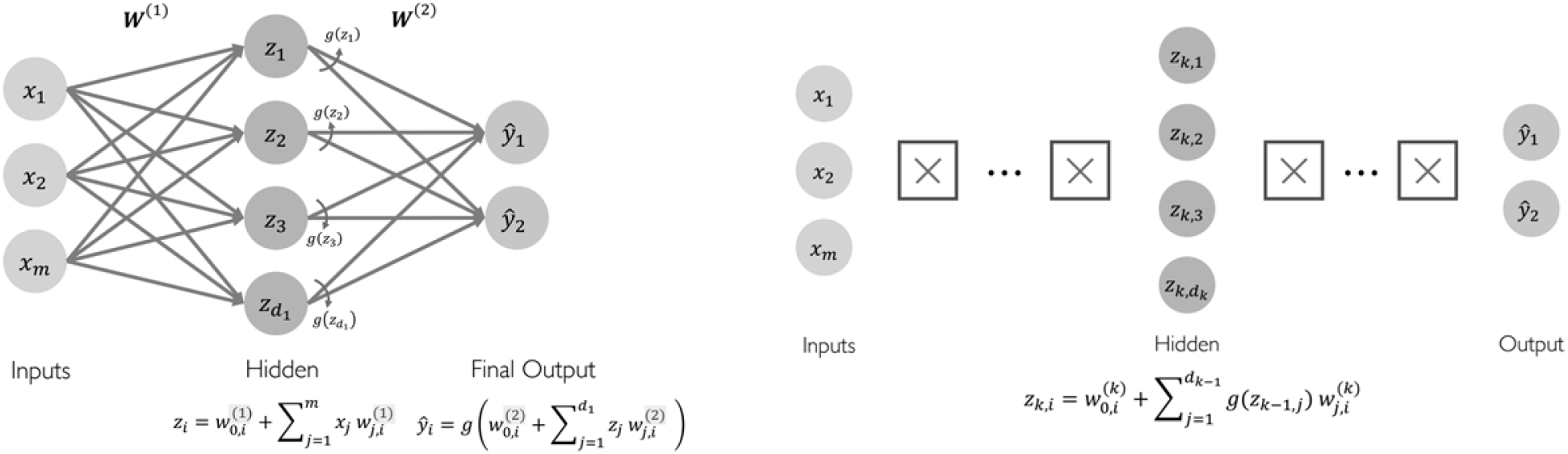
Single Layer Neural Network (left) and Deep Neural Network (right), where the value in the hidden layer *z* equals the weighted sums of the inputs plus a bias term, which are transformed using a non-linear activation function *g*. From MIT (*33*).

Equation 1. Softmax function. Where *K* is the number of classes, **s(x)** is a vector containing the scores of each class for instance **x** and *σ***(s(x))**_*k*_ is the estimated probability that instance **x** belongs to class *k* given the scores of each class for that instance. From Géron (*26*).

Several arguments can be made for developing DL-SDMs as an alternative to MaxEnt-SDMs. Firstly, there is the clarity in the interpretation of network’s output. The output will be two probabilities for each location, a probability for that location of belonging to class 1: species occurs, and a probability of belonging to class 0: species does not occur. A second argument, more interesting from an ecological point of view, is the possibility of taking into account the presence of other species as environmental predictors in DL-SDMs. This allows for the direct inclusion of biotic interactions that is not possible in MaxEnt. Researchers now often use a two-tiered approach, first running MaxEnt, and then separately modelling the output including co-occurrence of other species (*34, 35*). Including biotic interactions considerably improves model performance (*35*). Finally, a further incentive for developing DL-SDMs is their scalability. In MaxEnt-SDMs each species to be modelled requires the selection of a separate set of appropriate and uncorrelated input variables (*36, 37*). Given appropriate model structure, DL-SDMs can take the same complete set of (correlated) input features for each species. Next to this, there is also the potential of multi-classification in DL-SDM in which the model outputs the probability of presence for all species in a single instance. This would increase scalability as the network weights only need to be trained once, rather than separately for each species. Furthermore, these pretrained weights could be transferred to a new species dataset and retrained, likely reaching an optimal solution faster than starting from the default random initialization.

### 1.2 Aims of the study

The aims of this exploratory research are to (1) provide a proof of concept of DL-SDM, (2) compare performance of DL-SDM to MaxEnt-SDM and (3) to provide recommendations on the large scale practical implementation of DL-SDMs.

### 1.3 Research questions

Based on the aims of the research, the following research questions were defined:

1. What input data types are required for a DL-SDM?
2. Which evaluation methods are most suitable for a DL-SDM?
3. What type of DL-SDM architecture yields the best output?
4. How does the DL-SDM output compare to MaxEnt-SDM output?
5. How can DL-SDMs be implemented on a large scale?

## 2 Methods

### 2.1 Software

All source code for this research was written using Jupyter Notebook (*38*), based on a Python 3.6 kernel (*39*). The code, together with the input and output data is publicly available via github and can be found at: https://github.com/naturalis/trait-geo-diverse-dl.

### 2.2 Data preparation

The research project was structured in three separate stages. Firstly, a pilot model was made utilizing the same input species, occurrences and environmental predictors as recently used by Hendrix & Vos to model the niches of the world’s ungulates with MaxEnt (*40*). This choice was made to allow for a qualitative visual comparison of the results of the DL-SDM with MaxEnt-SDM. In the second stage, the number of occurrences in the pilot model was extended. This stage was used to gain deeper insight in the number of occurrences required for credible modelling performance for DL-SDM and potential improvements through changes in model architecture. Finally, in the third stage additional environmental predictors were included to assess their impact on model performance and potential improvements in model performance by changing model architecture.

#### 2.2.1 Pilot study

We used the occurrence data of 154 ungulate species and raster datasets for 41 abiotic environmental predictors relating to climate, topography and soil characteristics from the online repository of Hendrix & Vos (*40*). The occurrence data originated from the Global Biodiversity Information Facility (GBIF) website (*41*) and ranged between 10 - 882 observations per species (mean: 191 *±* 234 sd). The climatic raster data were sourced from the widely used BIOCLIM and ENVIREM datasets (*42, 43*). The soil characteristics rasters were sourced from the Land-Atmosphere Interaction Research Group, and topography rasters from the Harmonized World Soil database (*44*). All environmental rasters were transformed to a 5 minute spatial resolution. A full list of the variable descriptions of each raster can be found in Appendix A.

Starting with a csv file with filtered occurrences for a given species, the goal is to generate a dataframe including labeled positive and negative occurrence examples and the environmental variable values at these locations. This dataframe will form the input for the DNN. As no hard data on species absences exists, typically pseudo-absences are used instead (*45*). The steps taken to generate this dataframe are visualised in Figure 3 and detailed below. The code is provided in the *Stacking environmental rasters* and *Species and global prediction dataframes* notebooks in the pilot study folder in the repository. To generate pseudo-absences, circular buffers with 1000km radius were constructed around each occurrence point. These buffers were merged into a single ‘multipolygon’ shapefile. The environmental variable rasters were first stacked into a single multi-band raster and then clipped based on the extent of the multipolygon shapefile. A random selection of pseudo-absence locations was generated within the raster clip based on two constraints: (1) points were not allowed to be located within the sea and (2) points were not allowed to be located within raster cells with occurrences. For species with *<* 1000 occurrences, 1000 random locations were generated. For species with *>*1000 occurrences, the number of random locations was set equal to the number of occurrences. The resulting selection of pseudo-absence points and their longitude and latitude values were added to the csv file with filtered occurrences. Next, the environmental variable values for all locations were added to this dataframe. Each band in the stacked raster clip represented one of the 41 environmental variables. For all occurrence and pseudo-absence points, the cell number in which they were located was determined. By going iteratively through the raster bands, the cell values for all variables were extracted and added to the dataframe. The environmental variable values were scaled by subtracting the band mean and dividing by the standard deviation. This formed the dataset on which to train and test the DNN. To produce global predictions of species distributions after model training and testing, a separate dataset was made containing the scaled environmental variable values of all terrestrial cells in the stacked world raster map.

**Figure 2:**
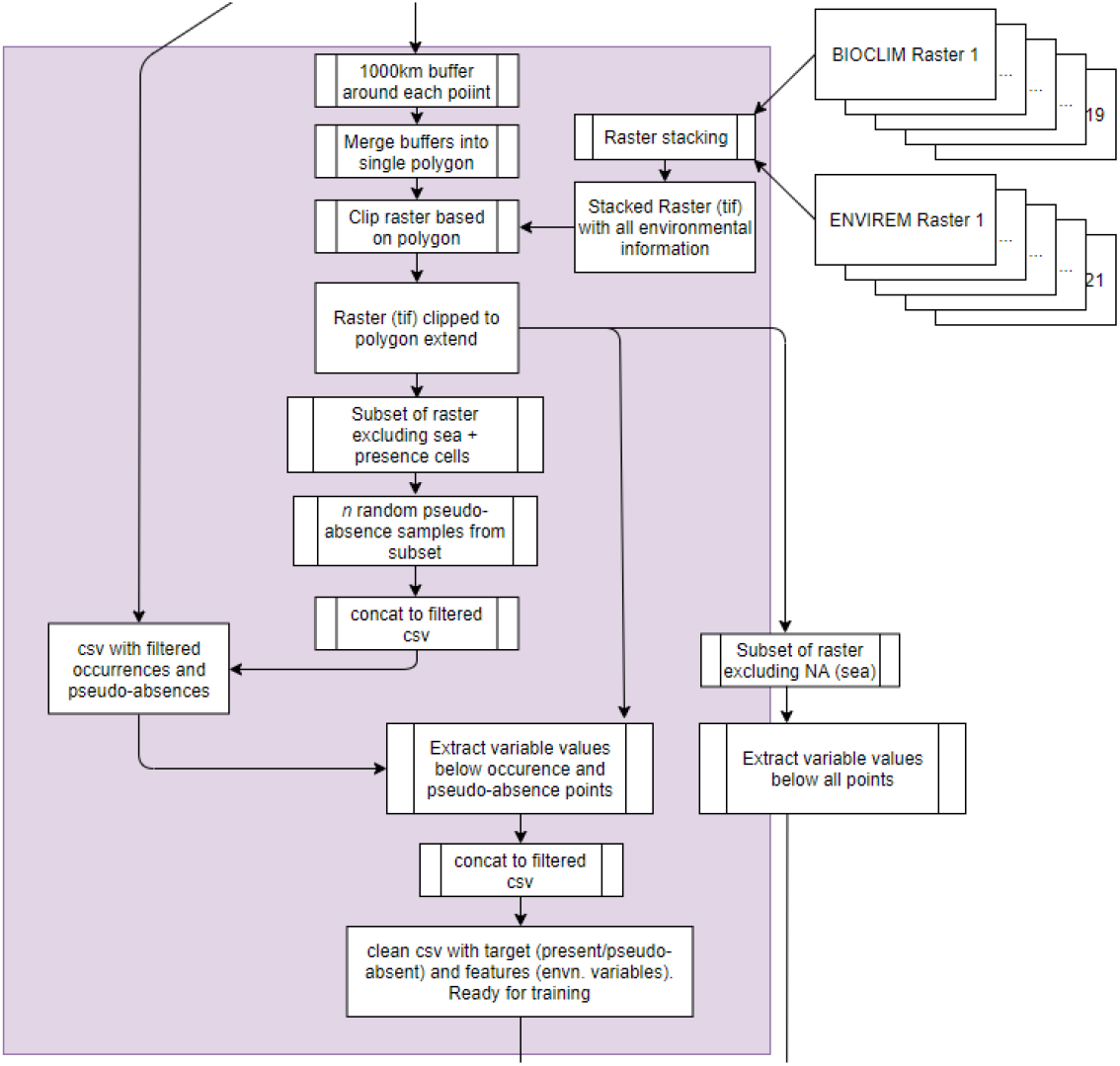
Workflow for preparing clean dataframe for DNN training and testing

**Figure 3:**
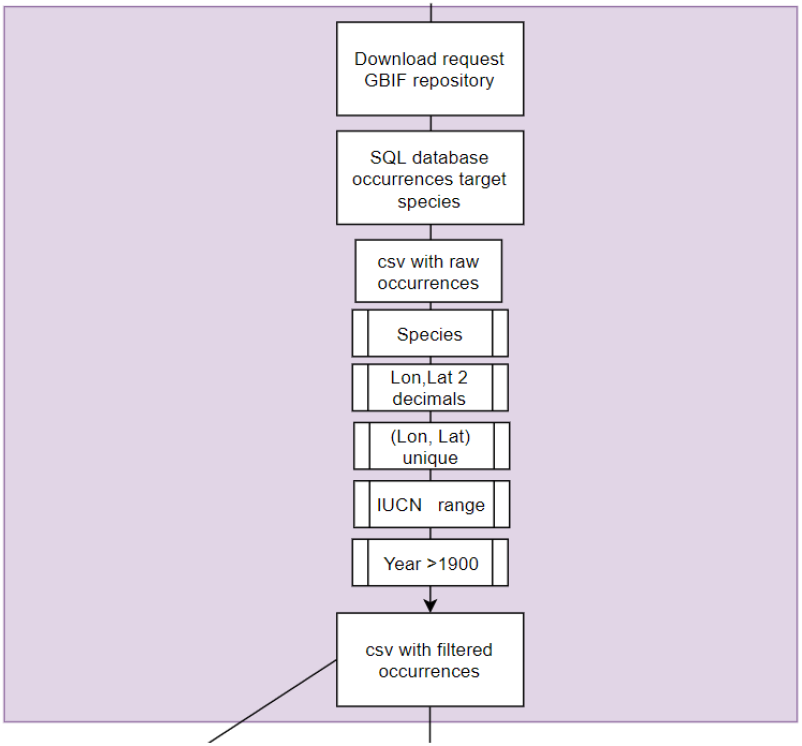
Workflow for filtering occurences from SQL database.

#### 2.2.2 Extended observations

In contrast to the pilot model, occurrence data was directly sourced from an SQL relational database containing all GBIF occurrences for the world’s ungulate species. These *raw* occurrences were first sorted on taxonomy and then filtered based on multiple criteria (Fig. 4), the code for which can be found in the *Filter GBIF records from SQL Database* notebook in the *data extended* folder in the repository. As a first filtering step, only occurrence records with at least two decimal values for longitude and latitude and records representing a unique longitude-latitude combination were included. Next, it was determined whether each occurrence was located within the species IUCN range by utilizing the publicly available IUCN species distribution range shapefiles (*46*). Finally, only records collected after 1900 and species with *>* 10 records after filtering were included. The total number of ungulate species included was 124, and the occurrences per species ranged between 10 - 58329 (mean: 1401 *±* 5798 sd). The subsequent process of generating pseudo-absences and extracting environmental values was the same as in the pilot project.

**Figure 4:**
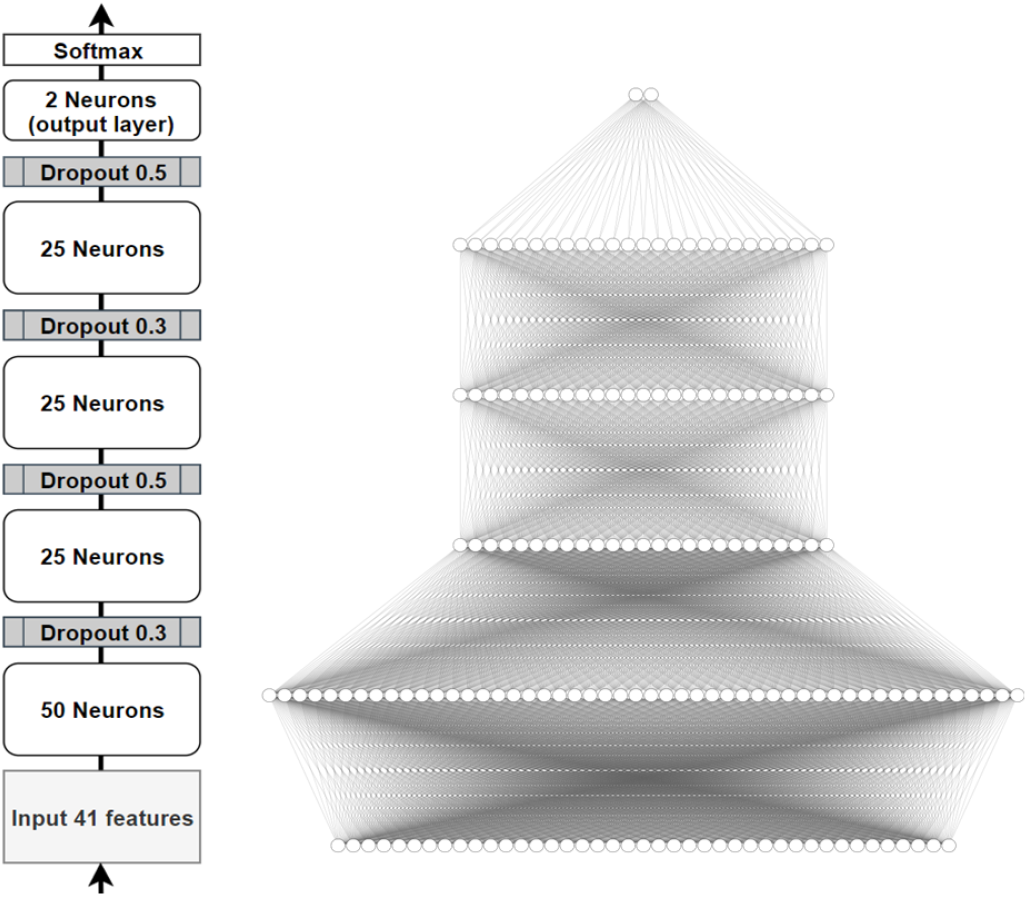
Left: pilot model architecture. Right: visualization of network connections in pilot model.

### 2.3.3 Extended observations and environmental variables

The same set of occurrence data was used as in the extended observation models. However, rather than generating pseudo-absence locations from within the buffers generated around each occurrence location, as in Hendrix & Vos (*40*), these were now sampled randomly from the entire world. This was done to increase the range of environmental variable values the model was exposed to during training on pseudo-absences and improve predictive capabilities at the global scale. For species with *>* 2000 occurrences, 2000 random locations were generated, and for species with more occurrences the number of random locations was set equal to the number of occurrences. Next to this, multiple biotic and abiotic variables were added to the environmental predictor dataset. These variables consisted of the occurrences of the other ungulate species in the dataset, as well as maps from the Atlas of World Conservation that represented: the world’s ecoregions, levels of human appropriation, human accessibility, habitat fragmentation, mammal species richness and plant species richness (*47, 48*). The code for processing, rasterizing and stacking these various additional environmental layers is listed in the *Environmental Raster Layers* notebook in the *data GIS extended* folder in the repository. The final stacked environmental raster contained 186 bands. A description for each variable is provided in Appendix B.

### 2.3 Data Analysis

#### 2.3.1 Model architecture and training

We first applied combinations between various learning rates, regularization functions, activation functions and optimization algorithms to a trial dataset of *Capreolus capreolus* to guide model construction (Table 1). The model structure was kept fixed with two hidden layers containing 50 and 25 neurons, a batch size of 100, and 500 epochs for training. In terms of model performance. we looked at the average loss, accuracy and AUC value for each of these hyperparameters (Table 2). The outcomes indicated the best performances were obtained using L2 or no regularization, a ReLu activation function, RMSProp or Adam optimization and a relatively high learning rate (0.001 or 0.0001). After this more systematic evaluation we attempted to further improve model performance by (1) adjusting the number of layers, (2) using drop-out as an alternative regularization method and (3) adjusting batch size and number of epochs. The final architecture of the pilot model consisted of four hidden layers with drop-out in between each layer (Fig. 5). We used Python’s Keras module to build the DNN (*49*) and trained the model using a batch size of 75 for 125 epochs, with a learning rate of 0.001 and using Adam optimization. As many datasets were imbalanced, with considerably more pseudo-absences than presence-locations, datasets were randomly shuffled, then split into training (85%) and test sets (15%) using a stratified approach. Furthermore, a balanced batch generator was used during training (*50*).

**Table 1:**
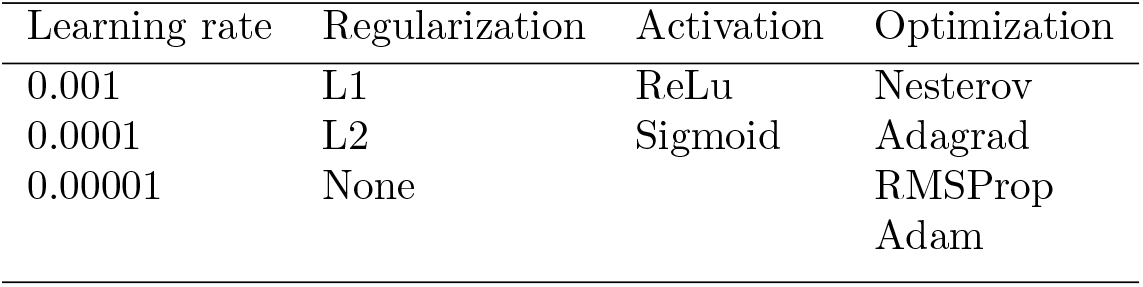
Combinations between multiple hyperparameters used to guide model construction (*n* = 360 runs). See glossary for definitions of hyperparameters.

| Learning rate | Regularization | Activation | Optimization |
| --- | --- | --- | --- |
| 0.001 | L1 | ReLu | Nesterov |
| 0.0001 | L2 | Sigmoid | Adagrad |
| 0.00001 | None |  | RMSProp |
|  |  |  | Adam |

**Table 2:** Model performance on *Capreolus capreolus* trial dataset with various hyperparameter combinations (mean *±* sd, *n* = 360 runs). See glossary for definitions of performance metrics.

| Hyperparameter | Average Loss | Average accuracy | Average AUC |
| --- | --- | --- | --- |
| L1 | $2.38 \pm 1.76$ | $0.65 \pm 0.13$ | $0.70 \pm 0.17$ |
| L2 | $0.89 \pm 0.38$ | $0.73 \pm 0.11$ | $0.81 \pm 0.11$ |
| None | $0.57 \pm 0.24$ | $0.74 \pm 0.12$ | $0.81 \pm 0.14$ |
| ReLu | $1.26 \pm 1.31$ | $0.77 \pm 0.10$ | $0.83 \pm 0.12$ |
| Sigmoid | $1.30 \pm 1.31$ | $0.65 \pm 0.13$ | $0.71 \pm 0.15$ |
| Nesterov | $1.46 \pm 1.43$ | $0.70 \pm 0.12$ | $0.78 \pm 0.12$ |
| Adagrad | $1.85 \pm 1.65$ | $0.63 \pm 0.14$ | $0.68 \pm 0.18$ |
| RMSProp | $0.90 \pm 0.89$ | $0.76 \pm 0.11$ | $0.82 \pm 0.12$ |
| Adam | $0.91 \pm 0.86$ | $0.75 \pm 0.11$ | $0.81 \pm 0.13$ |
| LR:0.001 | $0.67 \pm 0.34$ | $0.77 \pm 0.11$ | $0.82 \pm 0.14$ |
| LR:0.0001 | $1.19 \pm 1.24$ | $0.72 \pm 0.12$ | $0.79 \pm 0.13$ |
| LR:0.00001 | $1.98 \pm 1.62$ | $0.64 \pm 0.13$ | $0.71 \pm 0.16$ |

**Figure 5:**
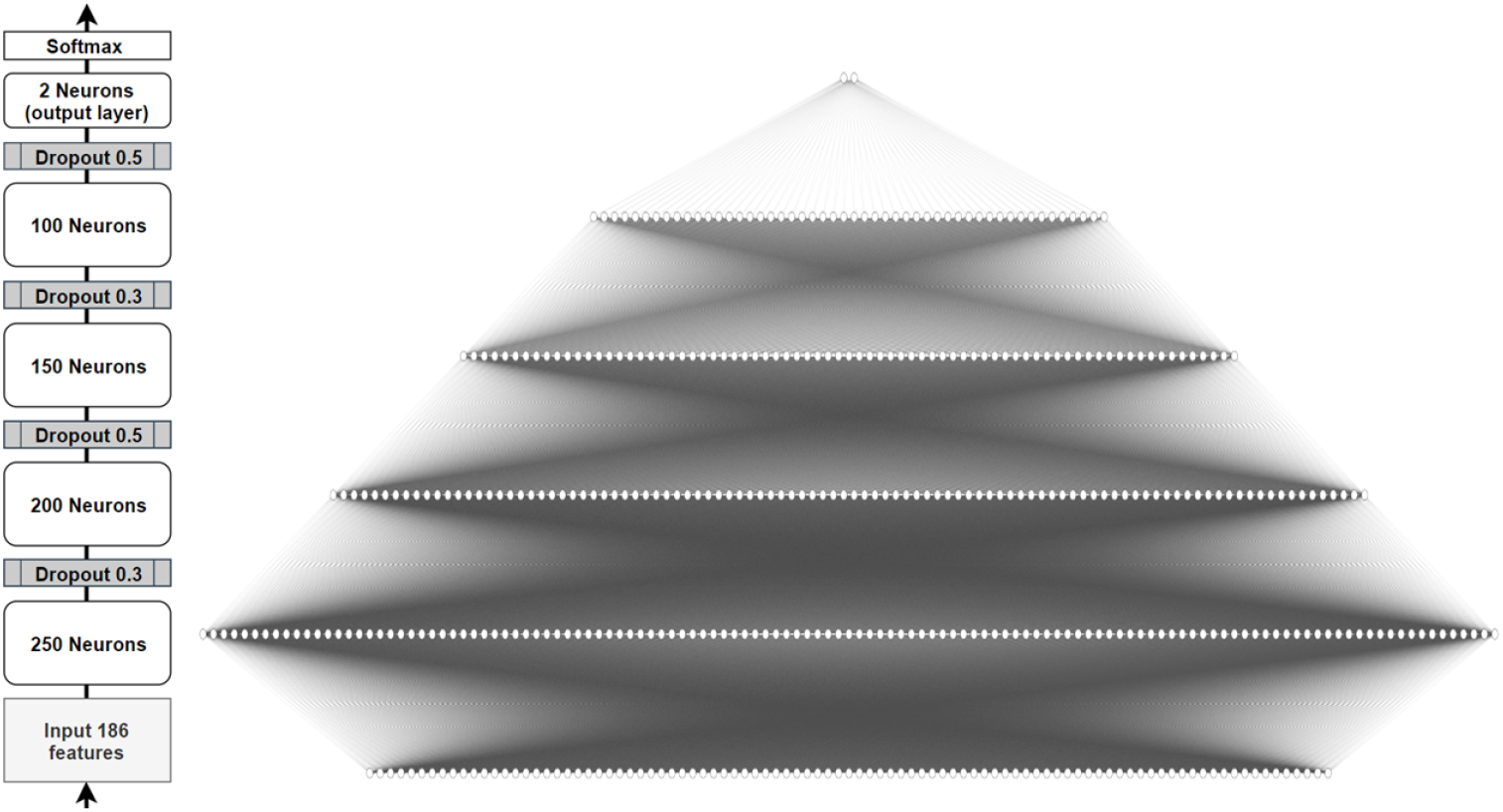
Left: model architecture extended observations and variables. Right: visualization of network connections in the model, the actual model contained twice as many neurons in the input and hidden layers.

The architecture for the model with extended observations was kept the same as the pilot model, as adding layers or altering drop-out rates did not seem to improve performance in the trial dataset. For the model with extended observations and variables, the number of layers and drop-out rates was kept the same, but the number of neurons in each hidden layer was increased to 250, 200, 150 and 100 neurons respectively, and the number of epochs was increased to 250 (Fig. 6). The code for training the models can be found in the *Train_DNN* notebooks.

**Figure 6:**
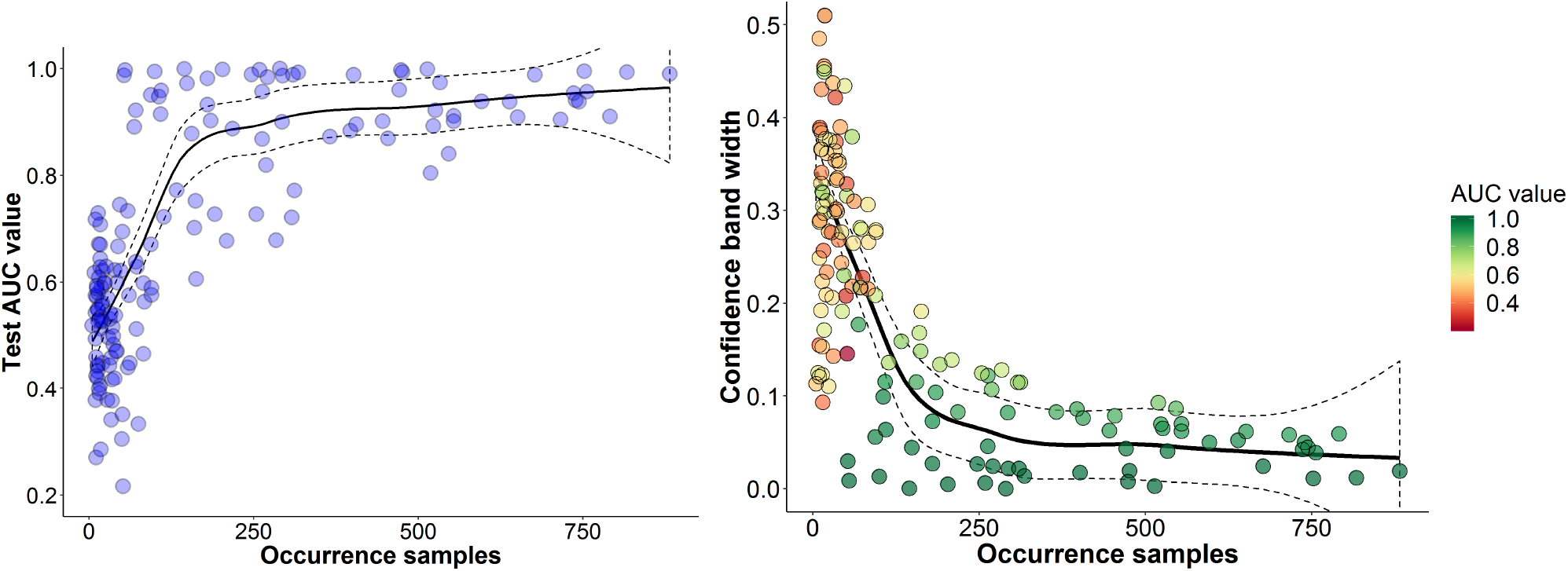
DNN pilot model performance. Left: change in model AUC with increasing occurrence samples. Right: Change in the width of the confidence interval around the model’s AUC value based on a bootstrapping procedure with 1000 repetitions. A LOESS smoother with a default span of 0.8 and 95% confidence intervals is fitted as a trendline in both graphs.

#### 2.3.2 Model evaluation

Evaluation of the models was the same for the pilot, extended observations, and extended observations and variable models. The DNN was run five times for each species. During each run, the test loss, accuracy and AUC value were stored and 95% lower and upper confidence bounds around the AUC value were estimated using a bootstrapping procedure with 1000 repetitions. The average test loss, accuracy, AUC and associated 95% confidence intervals over the five runs were written to a text file. The model weights of the run with the highest AUC value were saved as a. h5 file, to later reconstruct it for making predictions of the species global distribution. We used the DeepExplainer function from the SHAP package developed by Lundberg (*51*) to calculate feature importance by approximating Shapley values (*52, 53*). The approach computes the contribution of a target feature to a model prediction by rerunning the prediction using all possible non-target feature combinations and again for these combinations now including the target feature. It then takes the average difference in predicted outcomes. As DNNs’ fixed network structure means they cannot actually exclude a feature, excluded features take on a reference value instead (*54*).

## 3 Results

### 3.1 Pilot study

The DNN pilot model showed increasing performance and decreasing variation in performance with higher availability of occurrence samples (Fig 6). Compared to the MaxEnt model used by Hendrix & Vos (*40*), the pilot DNN model performed considerably worse when assessed over all species, with a large standard deviation, indicating high among species variation (Table 3). However, the difference in performance between the pilot DNN model and the MaxEnt model was relatively low if only species *>* 100 samples were taken into account. The predicted global distributions for a species with high, intermediate and low number of occurrence samples using both modelling approaches can be found in Figure 7. Associated variable importance for each of the individual models can be found in Appendix C.

**Table 3:**
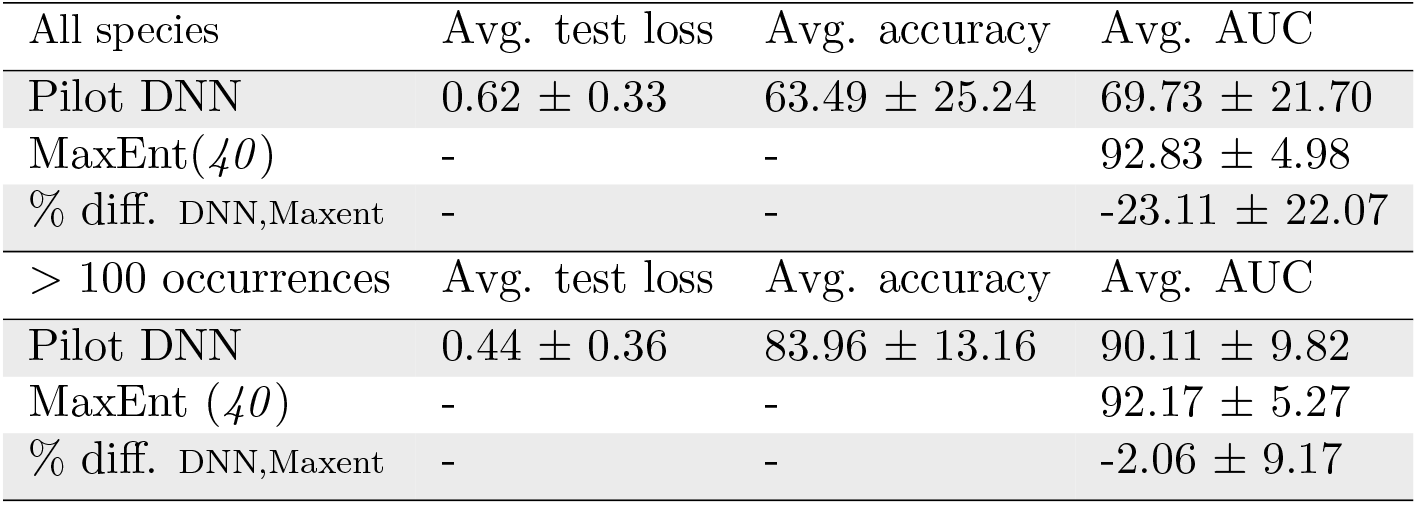
Model performance for all species (*n*=153) and subset of species *>* 100 occurrences (*n*=65) for Pilot DNN in comparison to MaxEnt study (*40*).

| All species | Avg. test loss | Avg. accuracy | Avg. AUC |
| --- | --- | --- | --- |
| Pilot DNN | $0.62 \pm 0.33$ | $63.49 \pm 25.24$ | $69.73 \pm 21.70$ |
| MaxEnt(40) | - | - | $92.83 \pm 4.98$ |
| % diff. DNN,Maxent | - | - | $-23.11 \pm 22.07$ |
| > 100 occurrences | Avg. test loss | Avg. accuracy | Avg. AUC |
| Pilot DNN | $0.44 \pm 0.36$ | $83.96 \pm 13.16$ | $90.11 \pm 9.82$ |
| MaxEnt (40) | - | - | $92.17 \pm 5.27$ |
| % diff. DNN,Maxent | - | - | $-2.06 \pm 9.17$ |

**Figure 7:**
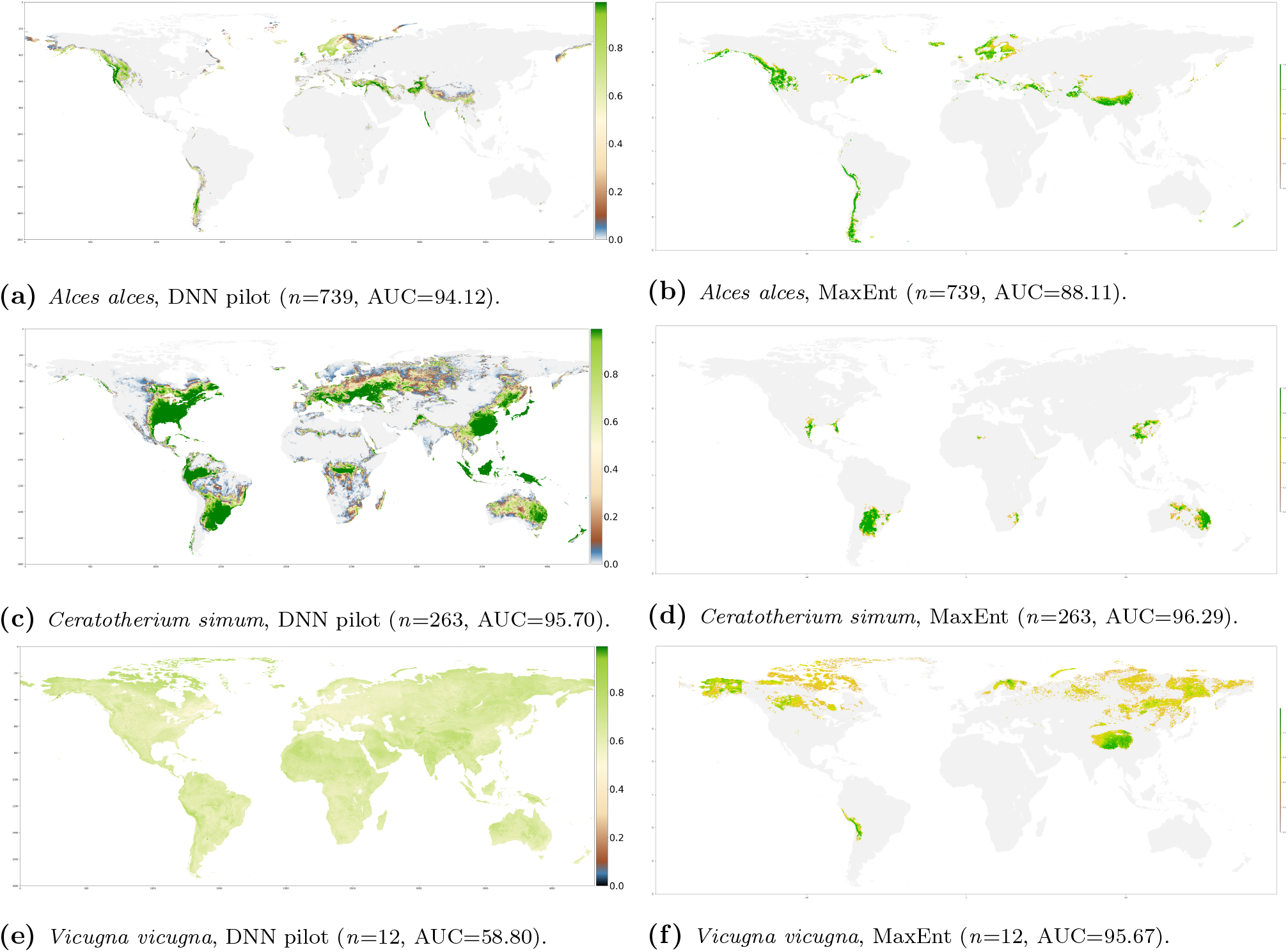
Global prediction maps for species with high, intermediate and low sample size compared between DNN pilot (left) and MaxEnt (right) SDM.

### 3.2 Extended observations

Including additional observations had mixed effects on model performance. Only for species with ≳ 500 observations, was there a clear improvement in terms of reduced test loss and increased AUC values (Table 4, Fig.8). This is also reflected in the changes in the predicted global distributions (Fig.10*._a,c,e_*). There was a large restriction in the predicted distribution of *Alces alces*, with 9966 occurrences, but not for the *Ceratotherium simum* and *Vicugna vicugna*, despite increases from 263 to 418 and from 12 to 61 occurrences respectively.

**Figure 8:**
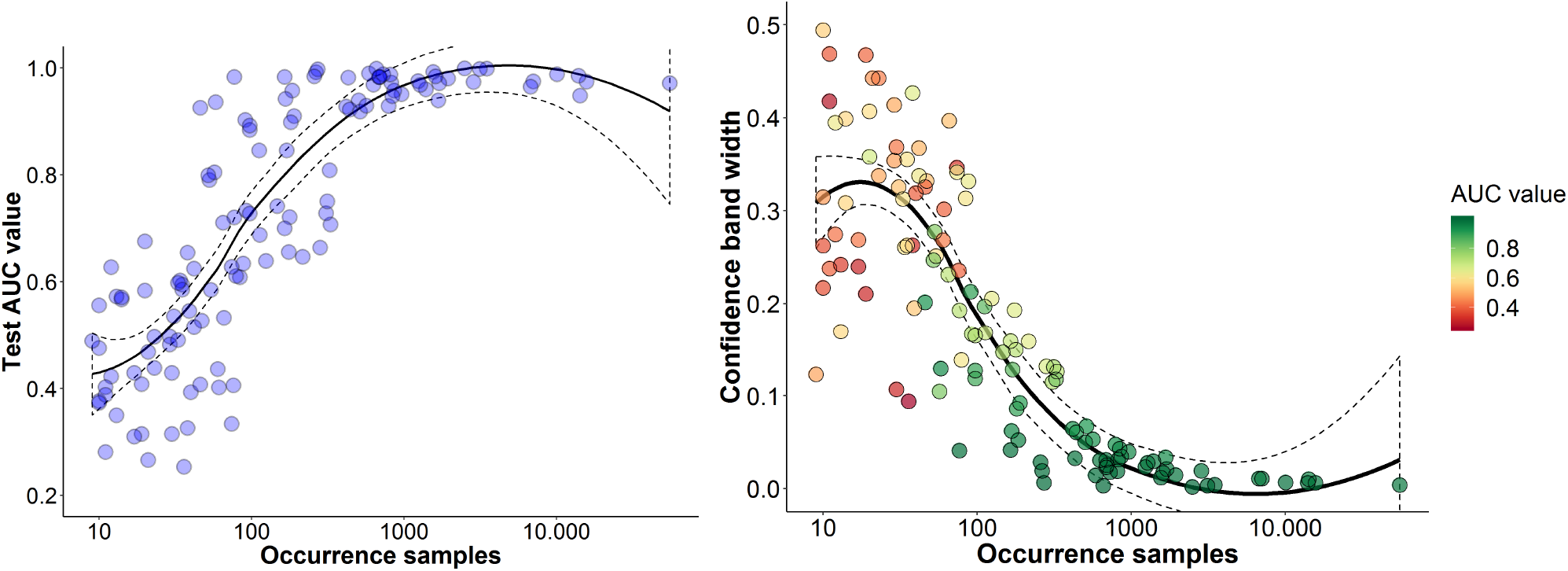
DNN extended observations model performance. Left: change in model AUC with increasing occurrence samples. Right: Change in the width of the confidence interval around the model’s AUC value based on a bootstrapping procedure with 1000 repetitions. A LOESS smoother with a default span of 0.8 and 95% confidence intervals is fitted as a trendline in both graphs.

**Table 4:** Model performance for all species (*n*=120) and subset of species *>* 500 occurrences (*n*=19) for Extended observations model in comparison to Pilot model.

| All species | Avg. test loss | Avg. accuracy | Avg. AUC |
| --- | --- | --- | --- |
| Extended obs. model | $0.58 \pm 0.35$ | $66.70 \pm 25.88$ | $71.91 \pm 23.42$ |
| Pilot model | $0.60 \pm 0.34$ | $65.88 \pm 25.34$ | $71.75 \pm 22.38$ |
| % diff. Extended,Pilot | $-0.02 \pm 0.28$ | $0.82 \pm 20.50$ | $0.16 \pm 16.04$ |
| $> 500$ occurrences | Avg. test loss | Avg. accuracy | Avg. AUC |
| Extended model | $0.22 \pm 0.09$ | $91.57 \pm 4.05$ | $96.77 \pm 2.11$ |
| Pilot model | $0.33 \pm 0.23$ | $88.20 \pm 7.04$ | $93.37 \pm 5.15$ |
| % diff. Extended,Pilot | $-0.11 \pm 0.18$ | $3.36 \pm 4.40$ | $3.40 \pm 3.69$ |

### 3.3 Extended observations and environmental variables

The inclusion of additional environmental variables and sampling pseudo-absences globally reduced the variation in AUC values and associated confidence intervals for species with sample sizes between *∼* 100 and 500 samples compared to the extended model (Fig.9). For species with *>* 500 observations, model loss, accuracy and AUC scores were all improved and there was relatively low variation in these metrics between species (Table 5). Although there was still considerable variation in performance measured across all species, the performance and predicted distribution of several species with *<* 500 occurrences did improve considerably, as can be seen in Figure 10*._b,d,f_*.

**Figure 9:**
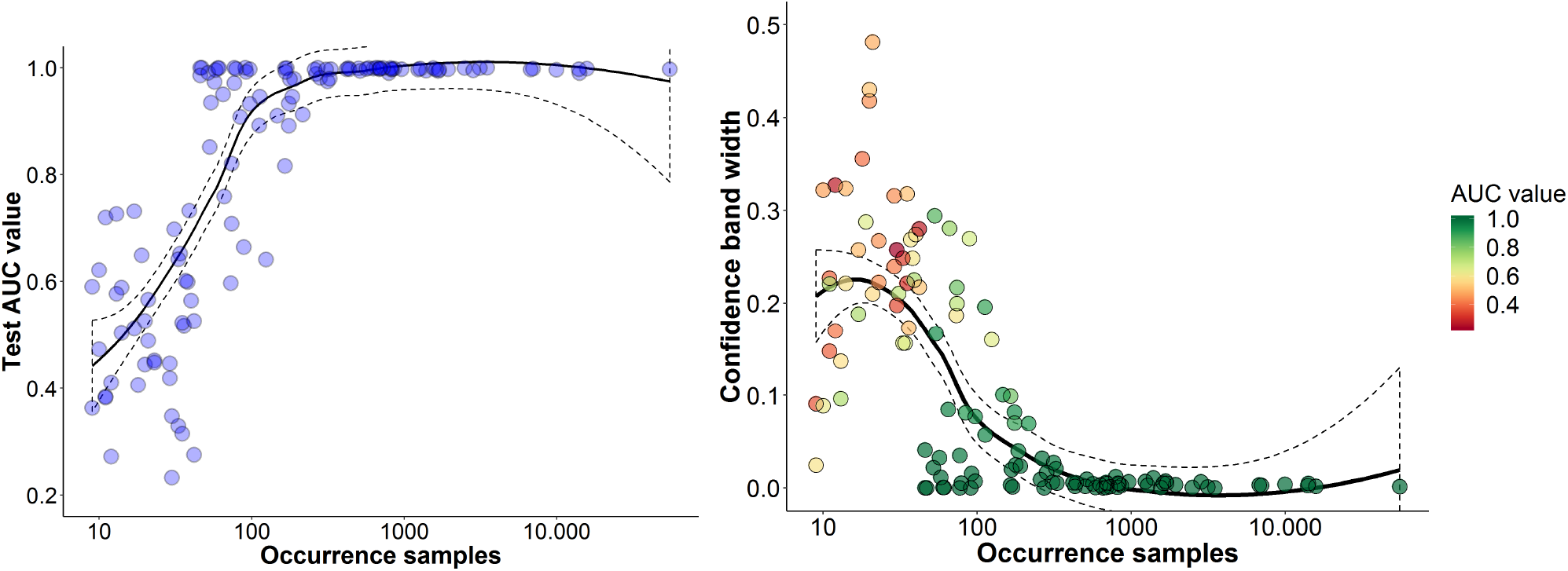
DNN extended observations and variables performance. Left: change in model AUC with increasing occurrence samples. Right: Change in the width of the confidence interval around the model’s AUC value based on a bootstrapping procedure with 1000 repetitions. A LOESS smoother with a default span of 0.8 and 95% confidence intervals is fitted as a trendline in both graphs.

**Table 5:** Model performance for all species (*n*=124) and subset of species *>* 500 occurrences (*n*=35) for Extended observations & variables model in comparison to Extended observations model.

| All species | Avg. test loss | Avg. accuracy | Avg. AUC |
| --- | --- | --- | --- |
| Extended ov model | $0.68 \pm 0.89$ | $76.10 \pm 24.32$ | $81.39 \pm 23.59$ |
| Extended model | $0.58 \pm 0.35$ | $66.99 \pm 25.71$ | $72.20 \pm 23.36$ |
| % diff. Extended ov, Extended | $0.11 \pm 0.74$ | $9.10 \pm 21.83$ | $9.19 \pm 16.22$ |
| $> 500$ occurrences | Avg. test loss | Avg. accuracy | Avg. AUC |
| Extended ov model | $0.10 \pm 0.06$ | $98.26 \pm 1.04$ | $99.74 \pm 0.24$ |
| Extended model | $0.21 \pm 0.09$ | $92.61 \pm 4.03$ | $97.06 \pm 2.12$ |
| % diff. Extended ov, Extended | $-0.11 \pm 0.07$ | $5.65 \pm 3.40$ | $2.69 \pm 1.98$ |

**Figure 10:**
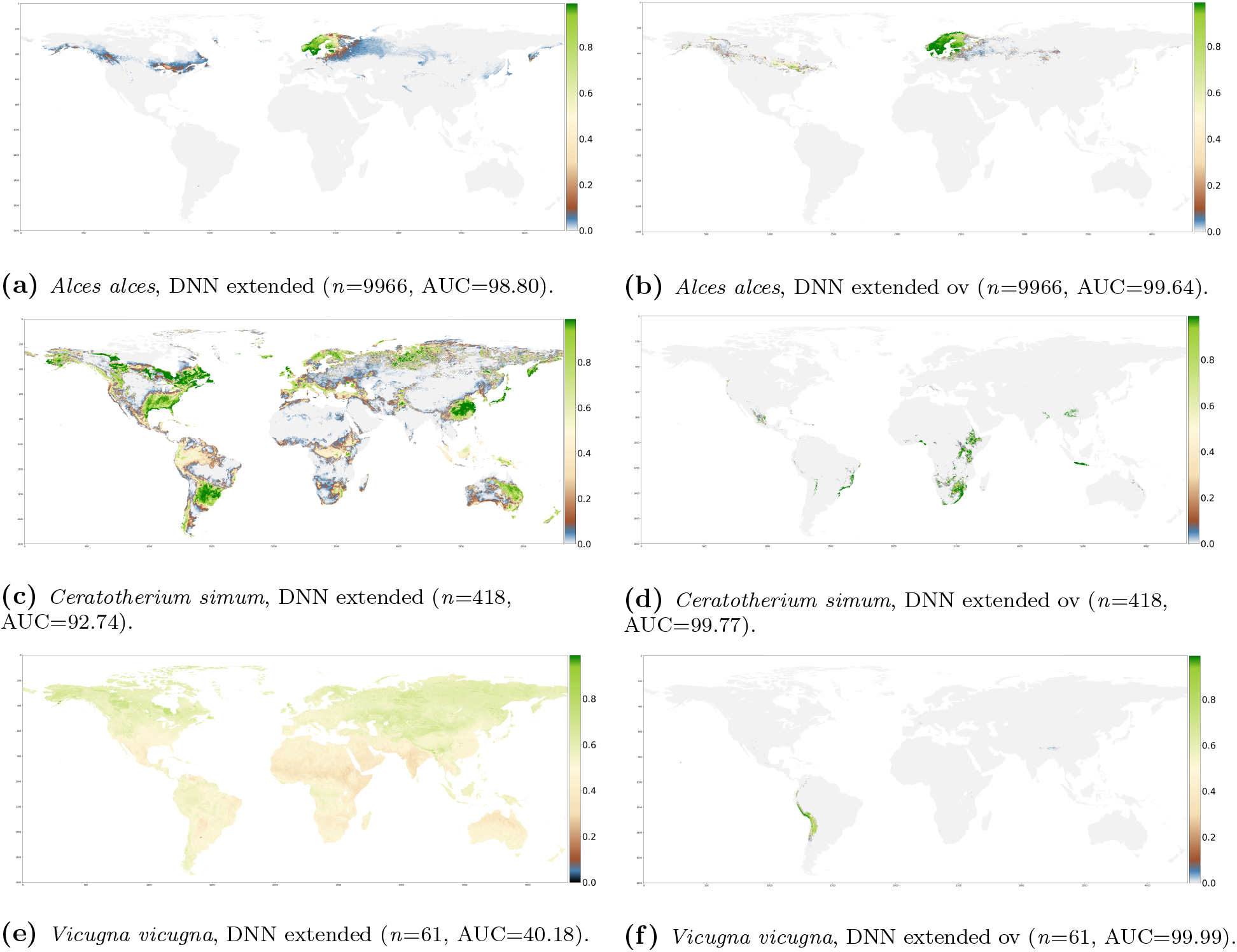
Global prediction maps for species with high, intermediate and low sample size compared between DNN extended observations (left) and DNN extended observations and variables (right) SDM.

Of the added environmental variables in the model, co-occurrence with another species was the most important feature for both *Alces alces* and *Ceratotherium simum* (Fig. 11). In the model for *Vicugna vicugna*, on the other hand, the most important features came from the same subset of abiotic variables as in the extended model, suggesting that the global pseudo-absence sampling strategy is responsible for the large improvement in model performance for this species. Three out of the five highest ranked features for *Ceratotherium simum* did not show a clear relationship between intermediate to high feature values and the impact on the model’s predicted probability of occurrence. Dependency plots indicated interaction effects occurring with other features (Fig. 12).

**Figure 11:**
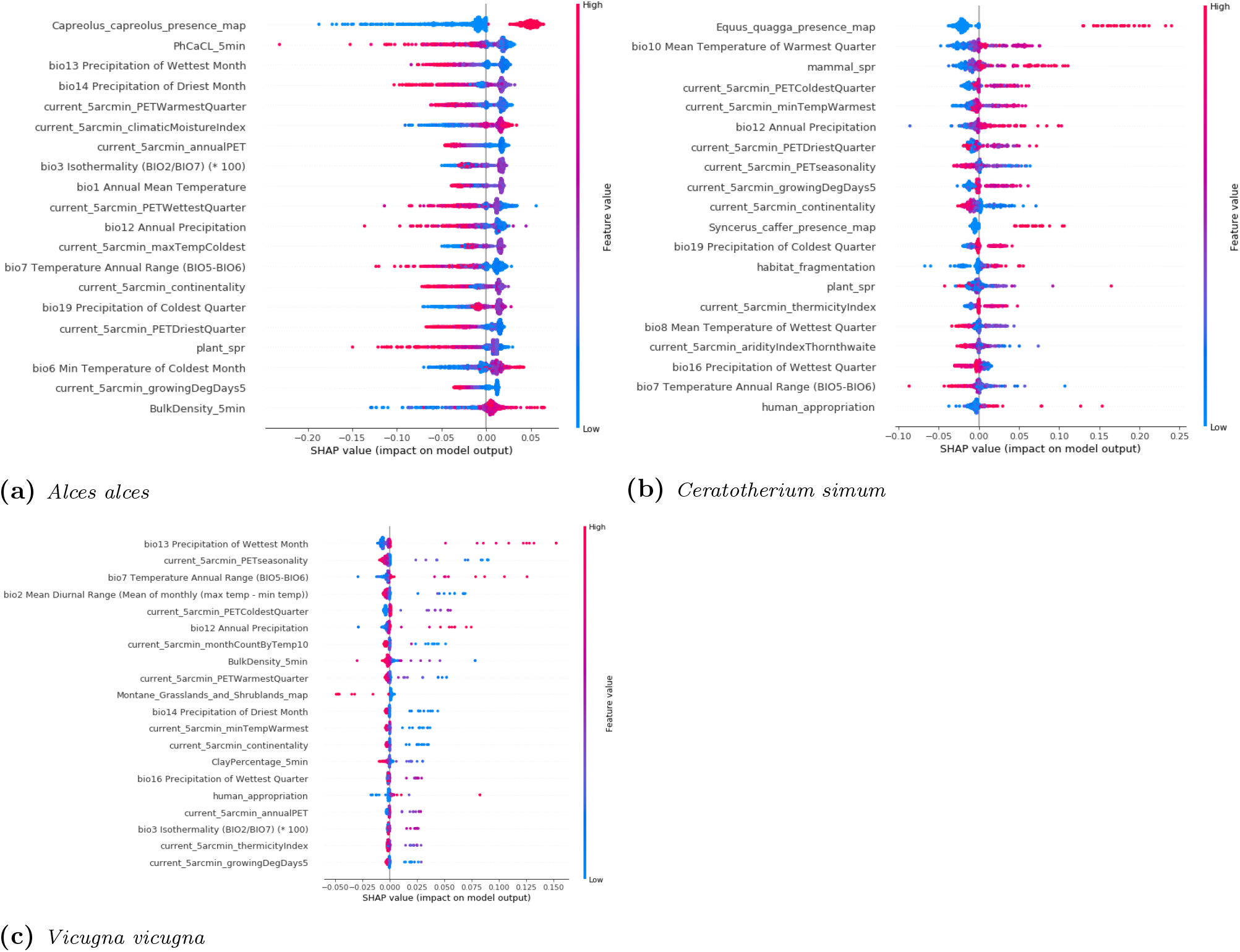
Feature importance for the extended ov models. Each dot represents an individual sample from the test dataset. Dot color indicates the value of the environmental predictor.The position on the x-axis indicates the impact on the predicted probability of occurrence.

**Figure 12:**
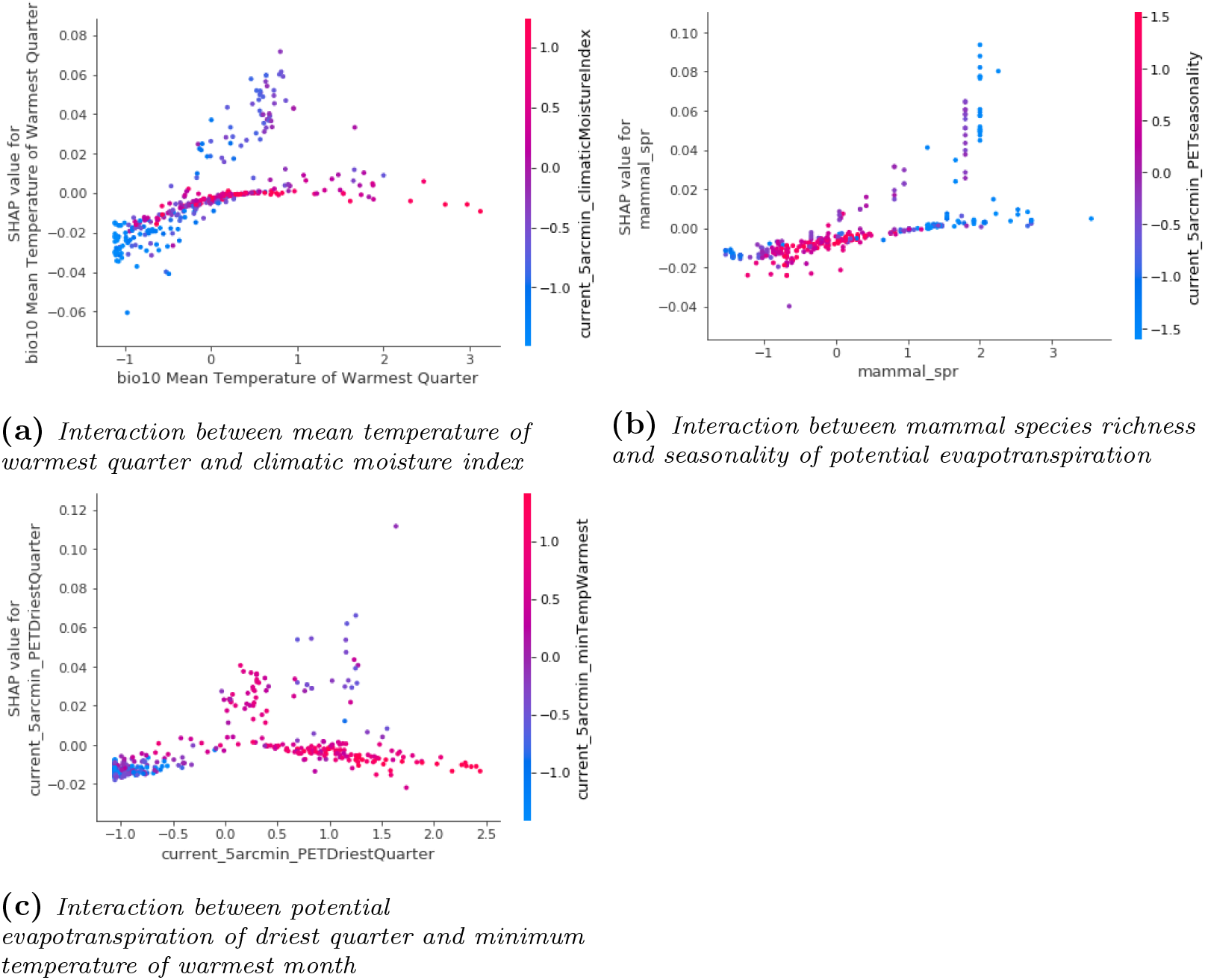
Feature dependencies for the *Ceratotherium simum* extended ov model. The impact of the target feature on the model’s predicted probability of occurrence is set against the target feature values. The second y-axis indicates the interaction effect occurring with a second feature.

The predicted probability of occurrence of *Ceratotherium simum* increases when going from low to intermediate temperature values in the warmest quarter and plateaus for high values. However, at intermediate levels (0 - 1), having conditions that are neither very arid nor moist (−1.0 - 0) increase the predicted probability of occurrence, whereas high moisture does not (Fig.12_*a*_). Reversing the scaling of the data shows that this corresponds to a combination of temperatures between around 21.1 - 39.3°C in the warmest quarter and moisture index levels of 53 - 55. Increased mammal species richness increases the predicted probability of occurrence. At high levels of mammal species richness (1.0 - 2.3; 94 - 227 spec.), having a low to intermediate seasonality in potential evapotranspiration (−1.5 - 0; 5.9 - 73.0 mm) further increases the predicted probability of occurrence (Fig.12_*b*_). Finally, both relatively low and high potential evapotranspiration in the driest quarter lower the predicted probability of occurrence. In between (0.0 - 1.5; 0.0 - 240.6mm), having intermediate to high values for the minimum temperature of the warmest month (0.0 - 1.0; 13.9 - 22.2 °C) increased the predicted probability of occurrence (Fig.12_*c*_).

## 4 Discussion

In this research we aimed to apply a Deep Learning approach to Species Distribution Modelling (DL-SDM). We also compared its performance to the well-established MaxEnt SDM on a limited dataset of the world’s ungulates.

### Relationship DNN and MaxEnt

Although the mechanics are still the subject of active research and debate (*55, 56*), the internal processes within a DNN share a similarity with the MaxEnt approach in that information flowing through the network converges to a maximum entropy solution (*57, 58*). In MaxEnt, this solution can be described as the distribution that minimizes the distance from the uninformed prior distribution of the ‘background’ feature set, but maintains the maximum amount of information contained in the distribution of the target feature set, i.e. has the same feature characteristics (mean, variance) as the feature set associated with the occurrence samples (*20*). Research by Schwarz & Tishby shows that going successively through each layer in a DNN, there is a trade-off between compression, or efficient representation of the information contained in the input features, and maintaining the predictive capabilities of the network (*57*) (Appendix E.1). The generalization capacity derived from this process does not occur in single-layered networks, and might partly explain their poor performance in SDM (*30, 31*). However, these findings are still debated and the process was not observed in research by Saxe (*55*) in networks utilizing ReLu activation functions (Appendix E.2). This is the most commonly used type of activation function and was also used in the networks in this research.

### Model comparison

Our model comparison based on a limited dataset of the world’s ungulates showed the DNN model performing worse for species with low and intermediate sample size and similar for species with a large sample size. DNNs typically require a large amount of training data to achieve high performance, which is related to to the large amount of parameters that need to be optimized (*59*). In this respect, the MaxEnt model is much less complex and it might explain why it performs better for species with few occurrence samples. However, sample size was not the only important determinant. The results of the extended ov model showed the selection of pseudo-absences was responsible for the large improvement in the global predicted distribution of *Vicugna vicugna*. By sampling negative labels only from within the IUCN range of the species in the pilot and extended model, these overfitted on the peculiarities of environmental conditions and generalized poorly when exposed to different conditions in other regions of the world. This also shows that if ‘pseudo-absences’ are not selected appropriately, an evaluation metric like the AUC value can be misleading. Both the MaxEnt model of the *Vicugna vicugna* and the DNN extended model of *Ceratotherium simum* achieved a high AUC, but their global predicted distribution showed poor generalization.

### DL-SDM improvement

One way to improve performance of the network model on small species datasets, is to apply transfer learning (*60*). Rather than learning the network weights starting from some random initialization, it is often beneficial for small datasets to use an existing model whose weights were pretrained for a similar classification task. This model can then be retrained on the small dataset, starting from the pretrained weights (*61*). This is a strategy that is often utilized in image recognition studies (*62– 64*), where well-known existing networks trained on thousands or millions of images such as Alexnet are retrained on the limited dataset available in the study. For DL-SDM, this could mean first training the model on an ecologically similar species with a large sample size and then apply transfer learning to retrain it on the target species with a small sample size. Alternatively, a single, deeper multi-classification model could be created that outputs the probability of presence for all species in a single instance. This model would then still need to be trained using resampling strategies to increase performance for species with few occurrences (*65*).

### Modelling shifting distributions

The DL-SDM in the current research was used to predict the distribution of the world’s ungulate species based on occurrence samples that were collected between 1900 and the present. Recently, there has been an increasing interest in modelling how species distributions might shift in the future following climate change (*66–68*). This could be modelled in DL-SDM by exposing a pretrained version of the current model to an adjusted set of environmental data, but the model would not be able to include species co-occurrences, as the distribution of the other species would likely change as well. Whether this is problematic would depend on the organism being modelled, for many of the ungulates in this research co-occurrences were shown to be important features, whereas one might expect plant distribution to be modelled accurately using only abiotic features.

The multi-classification model suggested earlier, which takes the occurrences of all species as inputs and also outputs the predicted occurrences of each species, could provide an approximation in two steps. In the first step the pretrained model is exposed to a new feature set including the adjusted abiotic environmental conditions, but the same set of species occurrences. The result can be framed as “*the predicted distribution of species X if only climatic conditions change and the distribution of other species remains the same*”. In the second step this pretrained model is then exposed again to the feature set with changed climatic conditions, but the species occurrences are replaced with the newly predicted distributions of all species. However, as the new distributions have arisen from a static process, which assumed the distribution of all other species remained the same, this would still provide a very rough approximation.

An alternative solution would be to create a dynamic version of the multi-classification model in the form of a Recurrent Neural Network (RNN) (*27*). RNNs are a type of neural network suitable for modelling sequential data. They have been very successful in language processing (*69, 70*), but are also used to approximate dynamic processes in climate modelling in computationally efficient ways (*71*). In an ecological setting, Lee & Donghyun (*72*) created RNN models to predict algal blooms in South Korean river systems. For DL-SDM, a dataset could be created that starts from current abiotic conditions, where at each time-step conditions are slightly changed in line with a certain climate scenario until reaching the predicted conditions in, for example, 2050. The co-occurrence features should be updated during each time step. At the first time step the current distributions of all species are used. The model then outputs the newly predicted distribution for all species under this small change in environmental conditions. In the next step, the co-occurrence feature values should be replaced by the newly predicted distribution values and so on until the end of the sequence. A potential downside to this approach would be that the RNN would initially have to be trained on a historic time-series dataset as well. This requires an explicit temporal link between the occurrence samples and environmental feature values that might be difficult to establish.

### Other applications

Another potential application of DL-SDMs is to combine them with image recognition techniques for automated species identification. Tools based on Convolutional Neural Networks (CNNs) are being developed to aid in species identification both in the field (*73*) and in museum collections (*74*). DL-SDM could provide an additional measure of certainty to a proposed identification by the CNN, by returning the probability that the species actually occurs at the locality the specimen was collected. If the process can be linked to a taxonomic relational database, another closely related species with a higher probability of occurrence at the specimen’s locality might then be proposed to the user.

## Conclusions

In this report we provided a proof of concept of DL-SDM using both a limited and an extended dataset of occurrences of the world’s ungulates. The required input consists of a selection of rasterized abiotic and biotic environmental predictor variables of the same spatial resolution. Notably, co-occurrences with other species proved an important environmental predictor for many of the ungulate species. Our final model required relatively few hidden layers (4) to gain good performance, in combination with a high number of neurons per layer (250, 200, 150, 100) and intermediate levels of drop-out regularization between layers (0.3, 0.5, 0.3, 0.5). Using a small dataset of the world’s ungulates and only abiotic predictors, the DL-SDM performed similar to the MaxEnt model of Hendrix & Vos for species with relatively high numbers of occurrences, and worse for species with low numbers of occurrences. Increasing the sample size, including species co-occurrences and improving pseudo-absence sampling resulted in large improvements in model performance and gave realistic distributions for species across a range of occurrence sample sizes. Implementing DL-SDMs on a larger scale will likely require the model to be transformed to a single multi-classification model.

## 4.1 Recommendations

To further explore the potential of DL-SDM, we recommend to (1) apply and adjust the current model to other groups of organisms, for example plants (2) to construct a single large multi-classification model for all species and compare it’s performance against the single species models in this research. (3) construct a temporal version of the model using a RNN framework to allow for modelling distribution shifts, for example under climate-change.

## 5 Acknowledgements

We would like to thank Elke Hendrix for discussing and clarifying the particularities of the limited ungulate dataset and MaxEnt model. Next to this, we would like to thank Dr. Anouschka Hoff from Wageningen University for the supervision provided to MR during the internship in which this work was produced.

## 7 Glossary

### 1. Accuracy

A metric for evaluating classification models, representing the fraction of predictions the model predicted correctly. In the binary classification example in this research, accuracy can be expressed as:

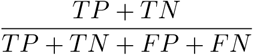

- *TP* = True Positive, a positive example correctly predicted positive.
- *TN* = True Negative, a negative example correctly predicted negative.
- *FP* = False Positive, a negative example falsely predicted as positive.
- *FN* = False Negative, a positive example falsely predicted as negative.

### 2. Area Under the ROC Curve (AUC)

The ROC curve plots the True Positive Rate (TPR) against the False Positive Rate (FPR) of the model predictions at different classification thresholds (*75*). The AUC value is a measure of the area under the ROC curve and indicates the quality of the model’s predictions integrated across all classification thresholds. Further definitions and examples are provided below.

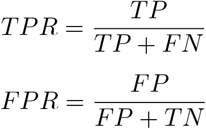

- *Classification threshold*. The model outputs the predicted probabilities of belonging to class 0 and 1. The classification threshold is the threshold value the output probability needs to have in order to be labeled as belonging to class 0 or class 1. Threshold values are problem dependent, i.e. how important the correct labeling of positive and negative examples is. A high threshold value might seem like a good option. However, increasing the threshold will both reduce false positive and true positive rates. That is, both negative examples with a relatively high output probability and positive examples with a relatively low output probability will not be labeled positive. This will therefore not be a good option if it is very important to correctly label all positive examples and there is a low cost to wrongly labeling negative examples.
- *ROC and AUC*. Example adapted from Park *et al*. (*76*). The figure shows four ROC curves. A perfect classification model (A) has an AUC value of 1. It always predicts the correct label. The diagonal line (D) represents a classification model no better at predicting labels than chance, it has an AUC value of 0.5. Between these extremes are curve B and C, with intermediate abilities to distinguish between different classes.

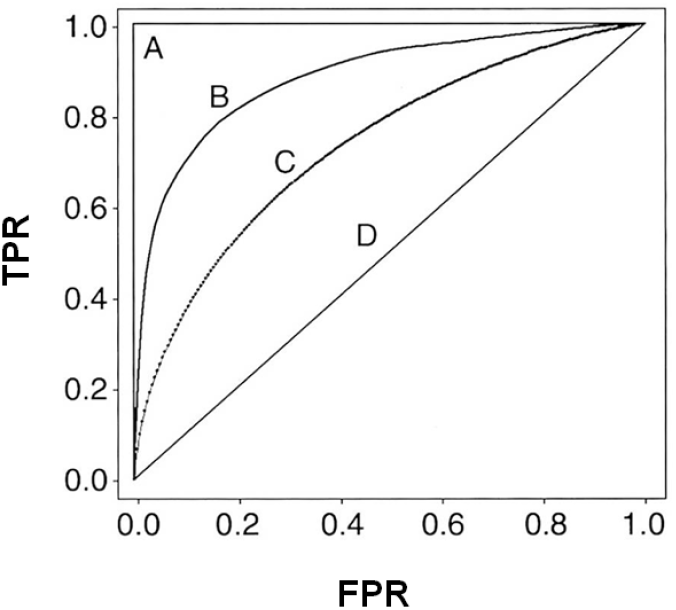

### 3. Batch size

The number of training examples the model works through before updating its internal parameters. In the case of DNNs, this refers to the updating of the network weights.

- *Balanced batch generator*. In an imbalanced dataset there are more samples of certain classes than others. This can have a negative effect on the predictive capabilities of a model, as during training it can be exposed to many batches that do not contain samples from the minority class. The model will still learn to achieve a high accuracy, but does this by simply classifying each sample as belonging to the majority class. A balanced batch generator resamples the dataset, usually by undersampling the majority class, to reduce the class imbalance in the batches passed to the model during training (*50*).

### 4. Epochs

The number of complete passes the model makes through the entire training dataset.

### 5. Learning rate

First read loss function. Optimization algorithms find the optimal set of parameter values required to minimize the loss function. As reviewing the change in model loss for each potential parameter value is inefficient, a step size is defined: the learning rate, which the optimizer uses to determine the next set of candidate weights to evaluate. As seen in the left figure below of a simplified loss landscape from Baughman & Liu (*77*), using a learning rate that is too low will result in very slow convergence and risks getting stuck in local minima, while a learning rate that is too high will overshoot the global minimum. The right figure from Li *et al*. (*78*) better illustrates the complex loss landscape of neural networks.

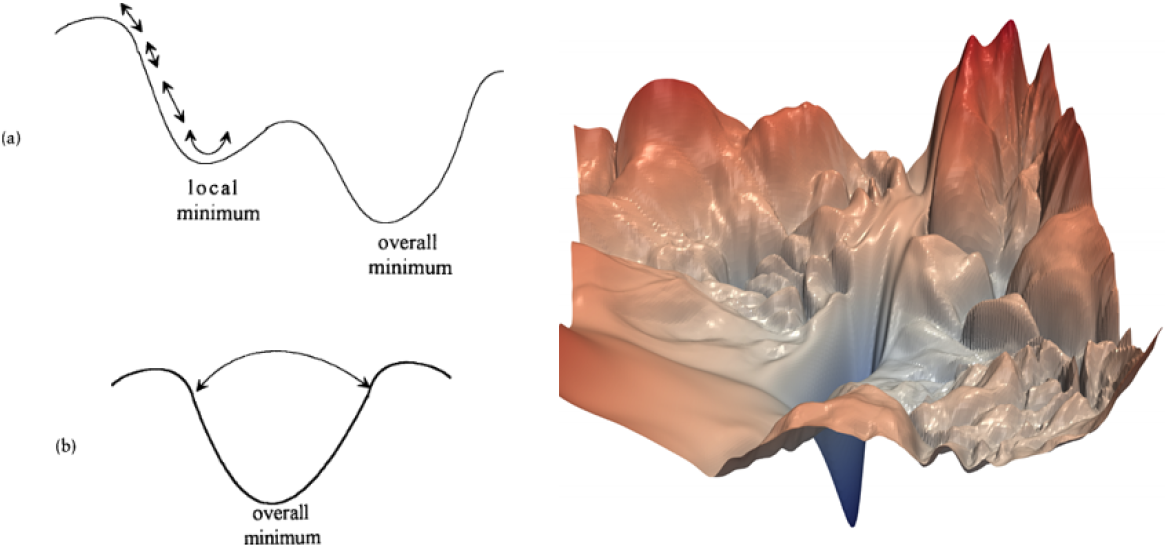

### 6. Loss function

In training the deep neural network, we are trying to minimize errors in classification. The loss function is used to evaluate the error value for a candidate set of network weights and bias terms identified by the optimization algorithm. An often used loss function for classification models is the cross-entropy function.

- *Cross entropy*. Measures the performance of a classification model that outputs a probability value between 0 and 1. If there are only two classes as in the current research, the cross-entropy function equals the log-loss function (*26*), expressed for a single training instance as:

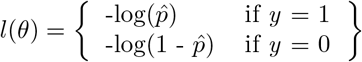

Where *l(θ)* is the estimated loss for the candidate set of weights and bias terms *θ*. The function is averaged over all training instances to estimate the cost function for the whole training set.

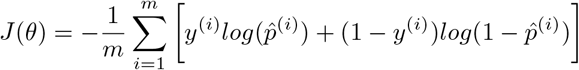 As explained by Géron (*26*), −log(x) increases as x aproaches 0 and decreases as x approaches 1. Therefore the cost will be high if the model estimates a probability close to 0 for a positive instance and low if it estimates a probability close to 1.

### 7. Non-linear activation function

Functions used to introduce non-linearity into the network. As each neuron in each layer computes a weighted sum of its inputs, the output of a network would remain a linear function, irrespective of how many layers are added, if no non-linear activation function is applied.

- *ReLU activation*. Short for Rectified Linear Unit, ReLU is the most commonly used activation function. It takes the weighted sum *z* of the inputs of each neuron. If *z* is equal to 0, the output of the neuron will be 0, if *z* is larger than 0, the output of the neuron is simply the weighted sum *z*, as seen in the figure from Sharma (*79*) below.

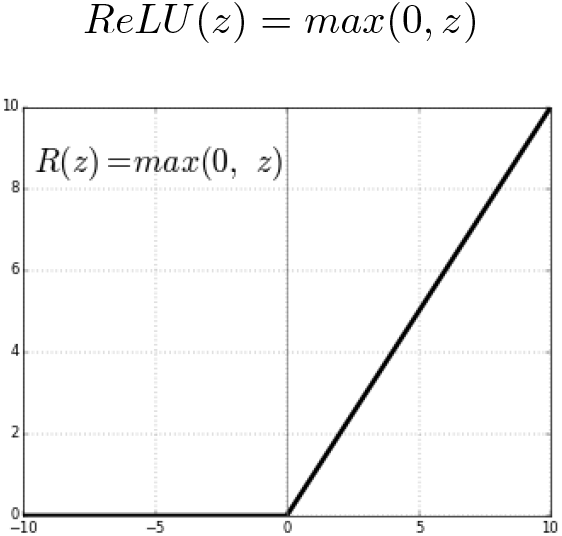
- *Sigmoid activation*. Long the default activation function for neural-networks, it has now started to fall out of favour to the ReLU function. This is because for deeper networks, there is a vanishing gradient problem illustrated in the figure below the equation, from Arunava (*80*). As the input values for the sigmoid function become larger or smaller, the derivative of the function becomes close to zero. Starting from the last layer in the network, optimization algorithms computes the gradient of the loss function for each parameter in the network and uses these to update the parameters based on the learning rate. This process is called backpropagation. If a sigmoid activation function is used, the gradients get increasingly small as the algorithm goes to the lower layers in the network, meaning the lower connection weights remain unchanged and the model cannot converge to a good solution (*26*).

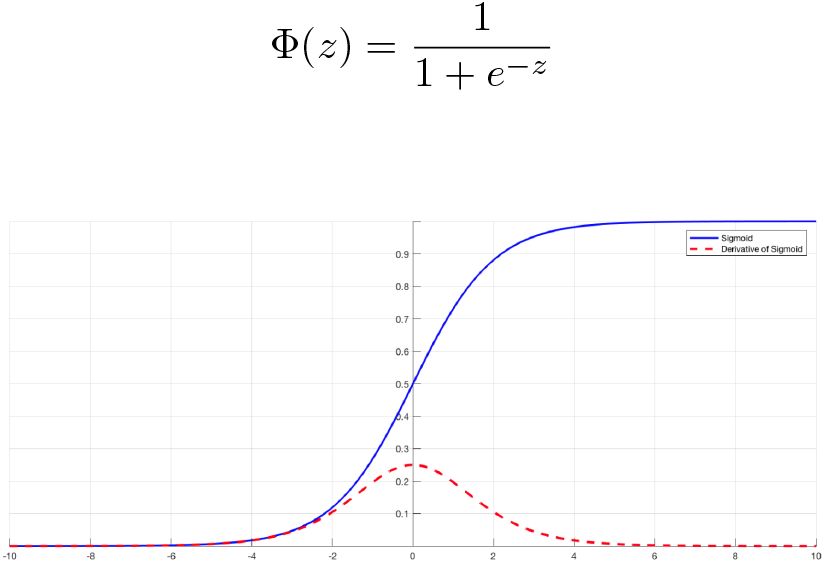

### 8. Optimization algorithms

Used to find the optimal set of parameter values (weights) in the network that minimize the loss function.

- *Gradient descent optimization*. The most common optimization algorithm used. Given a loss function *l* evaluated for a set of weights and bias terms *θ*, gradient descent adjusts *θ* using the following rule:

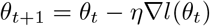 Where ∇*l* (*θ*) is the gradient of the loss function. This gives the direction in parameter space to increase the loss. Instead, gradient descent moves in the opposite direction (-∇*l* (*θ*)) based on the step size or learning rate *η* (*81*).
- *Gradient descent with momentum*. The Gradient descent algorithm was improved by including the concept of momentum, that can speed up movement along directions of strong improvement and better avoid local minima. This was achieved by introducing two additional parameters *v* and *µ* (*82, 83*). Where *v* is velocity, the exponential moving average of current and past gradients up to time step *t*, and *µ* is the momentum coefficient, between 0 and 1, that restricts the velocity. The updated rule becomes:

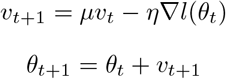
- *Nesterov optimization*. Variant of Gradient Descent with momentum that can speed up training and improve convergence. It measures the gradient of the cost function slightly ahead in the direction of the momentum (*26*). Notation wise, this difference is expressed in the update of the velocity vector *v* (*83*):

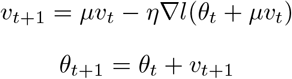
- *Adagrad optimization*. An optimizer providing an adaptive learning rate. Whereas the previous optimization algorithms used a single learning rate *η* for the set of parameters *θ*, Adagrad uses different learning rates for every parameter at every time step (*84*). The update rule for a single parameter can be expressed as:

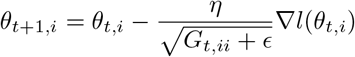 *G_t,ii_*, is a matrix containing the sum of the squares of the gradients of parameter *θ_i_* up to the current time step and *ϵ* is a smoothing term, typically 10^*−*10^, to prevent division by zero (*85*). For example, if three steps have been taken so far for parameter *θ*_1_, then the notation becomes:

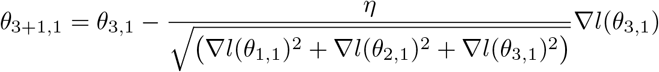 As with increasing time steps the sum of the gradients in the denominator also increases, the learning can become infinitesimally small over time and stop before the global optimum is reached (*85*).
- *RMSProp optimization*. Designed to handle the problem of Adagrad’s increasingly small learning rates. It only accumulates the gradients from the most recent iterations. In the expression, the diagonal matrix *G*_*t*_ is replaced by an exponentially decaying average over the past squared gradients (*85*). The parameter *β* is the decay rate, often set to 0.9 (*26*).

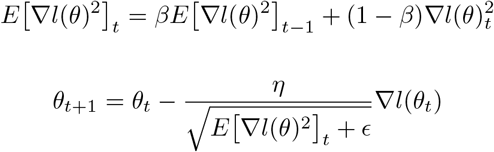
- *Adam optimization*. An optimizer combining the concepts of momentum and RMSProp. It stores an exponentially decaying average of both past gradients *m*_*t*_ and of past squared gradients *v*_*t*_ (*85, 86*).

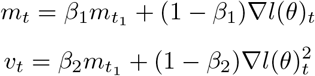 *β*_1_ represents the momentum decay and is usually set to 0.9, whereas the scaling decay *β*_2_ is usually set to 0.99 (*26*). However, with these values *v*_*t*_ and *m*_*t*_ are biased towards zero during the first few time steps (*86*). Therefore bias corrected values are used instead in the Adam update rule:

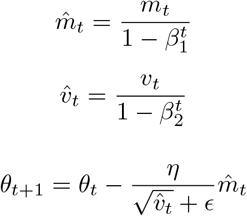

### 9. Regularization

Introduces a penalty term in the model’s loss function that penalizes model complexity to prevent overfitting.

- *L1 regularization*. Regularization term encouraging feature sparsity by setting the weights of the least important features to zero if parameter *α* is sufficiently large (*87*). The regularization part of the loss function can be expressed as:

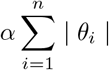
- *L2 regularization*. Regularization term that encourages weight values close to zero and the mean of the weights towards zero with a gaussian distribution. L2 regularization penalizes the squared values of the weights.

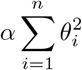
- *Drop-out regularization*. The approach randomly sets the activation of a collection of neurons to zero during training, dropping all their connections in the network during a single pass through the network and weight updating (*88*). This prevents neurons in the network from over-specializing on a specific feature in the training dataset, which results in poor model generalization. Typically drop-out rates between 0.1 and 0.5 are used. An example representation can be seen in the figure below from MIT (*33*).

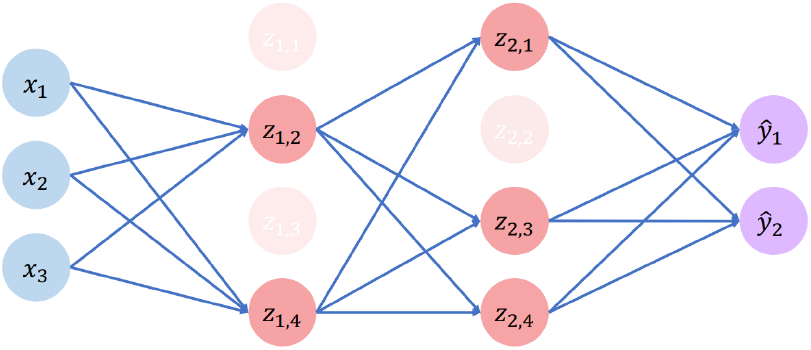

### 10. Shapley values

The approach has its origin in game theory and calculates the fair contribution of players to the outcome of a game with a collective pay-off (*52*).

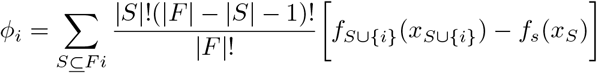

Where *φ_i_* is the contribution of player *i*. The order in which the players took turns can affect their contribution. To objectively assess the contribution of a single player therefore requires taking into account the possible sequences in which the game could have been played. This is addressed in the equation by using subsets, *S*, of the total set of players, *F*, excluding *i*. For each subset, the nominator in the equation takes all possible sequences, |*S*|!, in which the players could have played. If player *i* plays after this subset, then there are (|*F| − |S|* − 1) remaining players after *i* and there are (|*F| − |S|* − 1)! possible sequences in which they could have played. The total number of potential sequences considering all players that were in the game is given by |*F* |! in the denominator. Thus, the fraction in the equation represents the proportion of the total number of sequences accounted for by the subset. It functions as a weight for the outcome of the second part of the equation, which computes the difference in the pay-off between the subset including player *i* and excluding player *i* (*89*).

For determining variable importance, the equation is altered only slightly:

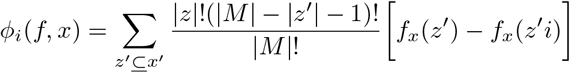

Where *M* is a vector of the full set of variables excluding the target feature, and *z’* is the variable vector with a subset of features set to a reference value (*53*).

#### A Environmental variables pilot model

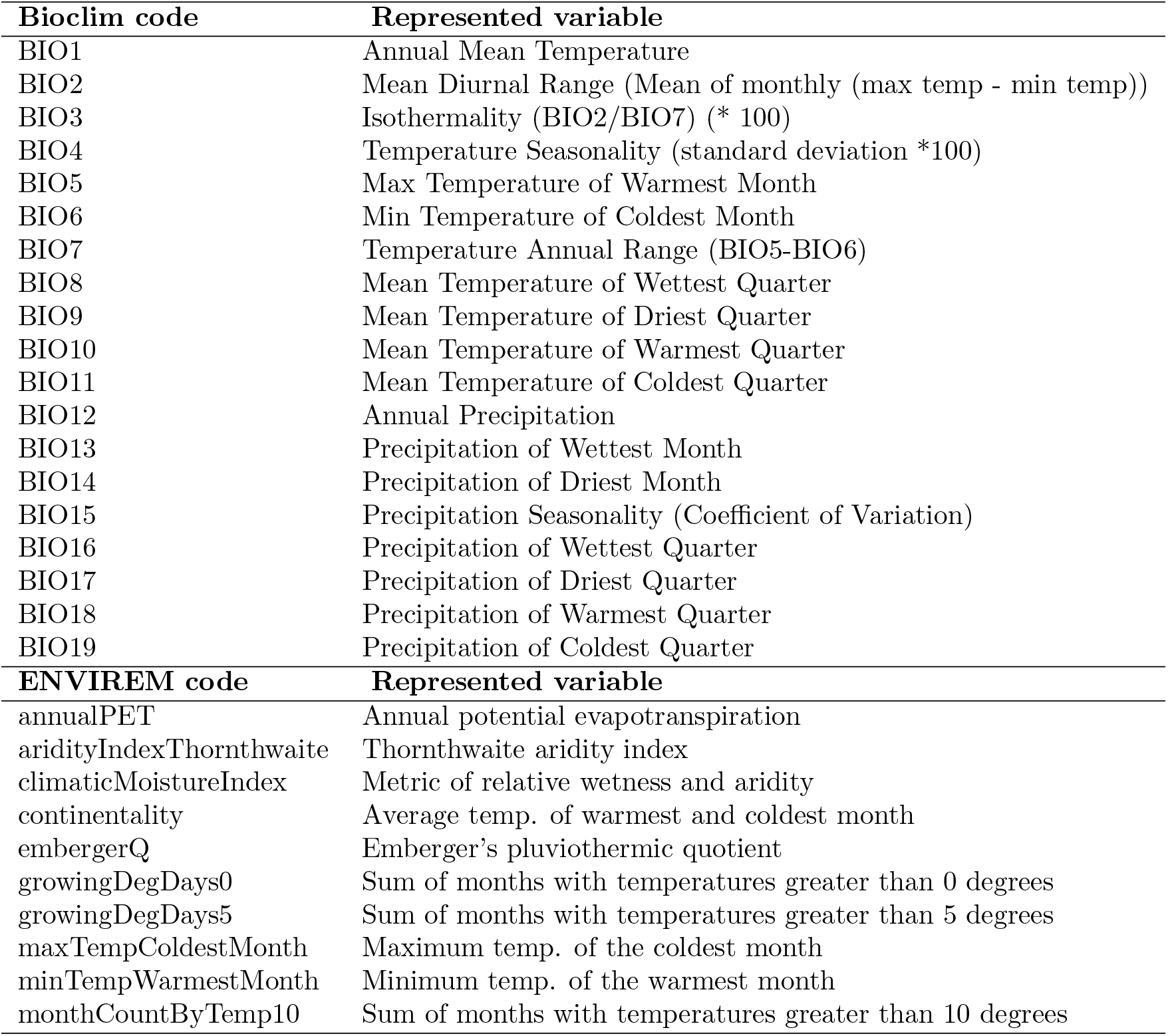

#### B Environmental variables extended ov model

| Ecoregion variables |  |  |
| --- | --- | --- |
| Boreal Forests and Taiga |  |  |
| Deserts and Xeric Shrublands |  |  |
| Flooded Grasslands and Savannas |  |  |
| Inland Water |  |  |
| Mangroves |  |  |
| Mediterranean Forests, Woodlands and Scrubs |  |  |
| Montane Grasslands and Shrublands |  |  |
| Rock and Ice |  |  |
| Temperature Broadleaf and Mixed Forests |  |  |
| Temperature Conifer Forests |  |  |
| Temperate Grasslands, Savannas and Shrublands |  |  |
| Tropical and Subtropical Coniferous Forests |  |  |
| Tropical and Subtropical Dry Broadleaf Forests |  |  |
| Tropical and Subtropical Grasslands, Savannas and Shrublands |  |  |
| Tropical and Subtropical Moist Broadleaf Forests |  |  |
| Tundra |  |  |
| Ecoregion attribute variables |  |  |
| Habitat fragmentation |  |  |
| Human accessibility |  |  |
| Human appropriation |  |  |
| Mammal species richness |  |  |
| Plant species richness |  |  |
| Species occurrence variables |  |  |
| Aepyceros melampus | Equus zebra | Pelea capreolus |
| Alcelaphus buselaphus | Eudorcas rufifrons | Phacochoerus aethiopicus |
| Alces alces | Eudorcas thomsonii | Phacochoerus africanus |
| Antidorcas marsupialis | Gazella bennettii | Philantomba maxwellii |
| Antilocapra americana | Gazella dorcas | Philantomba monticola |
| Antilope cervicapra | Gazella gazella | Potamochoerus larvatus |
| Axis axis | Gazella subgutturosa | Potamochoerus porcus |
| Axis porcinus | Giraffa camelopardalis | Procapra picticaudata |
| Bison bison | Hippopotamus amphibius | Pseudois nayaur |
| Bison bonasus | Hippotragus equinus | Pudu puda |
| Blastocerus dichotomus | Hippotragus niger | Rangifer tarandus |
| Bos javanicus | Hydropotes inermis | Raphicerus campestris |
| Boselaphus tragocamelus | Hyemoschus aquaticus | Raphicerus melanotis |
| Capra ibex | Hylochoerus meinertzhageni | Raphicerus sharpei |
| Capra nubiana | Kobus ellipsiprymnus | Redunca arundinum |
| Capra pyrenaica | Kobus kob | Redunca fulvorufula |
| Capra sibirica | Kobus leche | Redunca redunca |
| Capreolus capreolus | Kobus megaceros | Rhinoceros unicornis |
| Capreolus pygargus | Kobus vardonii | Rupicapra pyrenaica |
| Capricornis crispus | Litocranius walleri | Rupicapra rupicapra |
| Continued on next page |  |  |

| Species occurrence variables |  |  |
| --- | --- | --- |
| Capricornis swinhoei | Madoqua guentheri | Rusa unicolor |
| Catagonus wagneri | Madoqua kirkii | Sus barbatus |
| Cephalophus dorsalis | Mazama americana | Sus scrofa |
| Cephalophus jentinki | Mazama gouazoubira | Sylvicapra grimmia |
| Cephalophus natalensis | Mazama nana | Syncerus caffer |
| Cephalophus niger | Mazama pandora | Tapirus terrestris |
| Cephalophus nigrifrons | Mazama temama | Tayassu pecari |
| Cephalophus rufilatus | Muntiacus reevesi | Tragelaphus angasii |
| Cephalophus silvicultor | Nanger granti | Tragelaphus buxtoni |
| Cephalophus zebra | Nanger soemmerringii | Tragelaphus eurycerus |
| Ceratotherium simum | Neotragus batesi | Tragelaphus imberbis |
| Cervus elaphus | Neotragus pygmaeus | Tragelaphus oryx |
| Cervus nippon | Odocoileus hemionus | Tragelaphus scriptus |
| Connochaetes gnou | Odocoileus virginianus | Tragelaphus spekii |
| Connochaetes taurinus | Oreamnos americanus | Tragelaphus strepsiceros |
| Dama dama | Oreotragus oreotragus | Tragulius kanchil |
| Damaliscus lunatus | Oryx beisa | Tragulius napu |
| Damaliscus pygargus | Oryx gazella | Vicugna vicugna |
| Diceros bicornis | Ourebia ourebi |  |
| Equus grevyi | Ovibos moschatus |  |
| Equus hemionus | Ovis canadensis |  |
| Equus kiang | Ovis dalli |  |
| Equus quagga | Pecari tajacu |  |

#### C Feature importance

##### 1.

Feature importance of DNN pilot and MaxEnt models for top: *Alces alces*, centre:*Ceratotherium simum*, bottom: *Vicugna vicugna*.

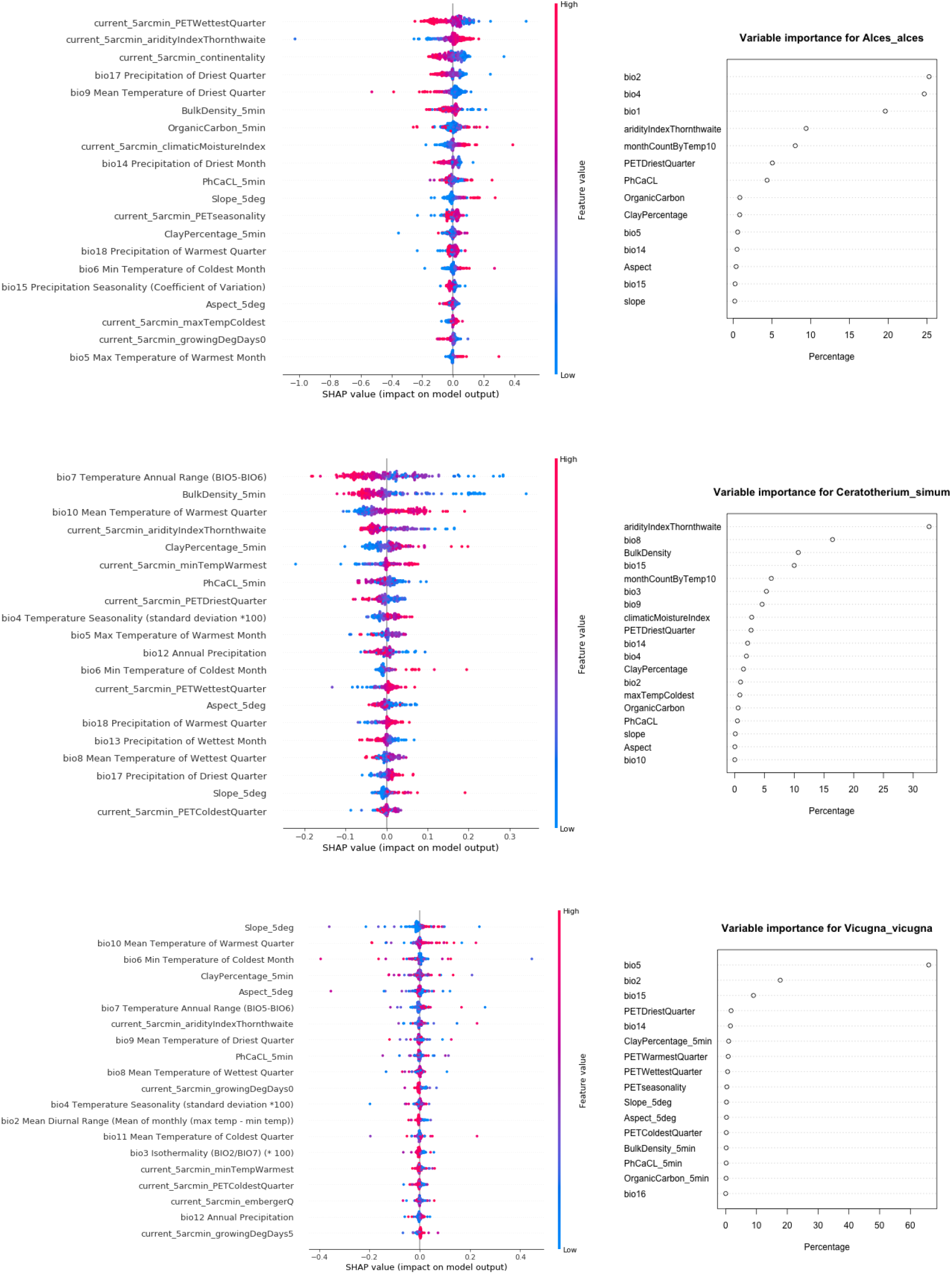

##### 2.

Feature importance of DNN extended model for top: *Alces alces*, centre: *Ceratotherium simum* and bottom: *Vicugna vicugna*.

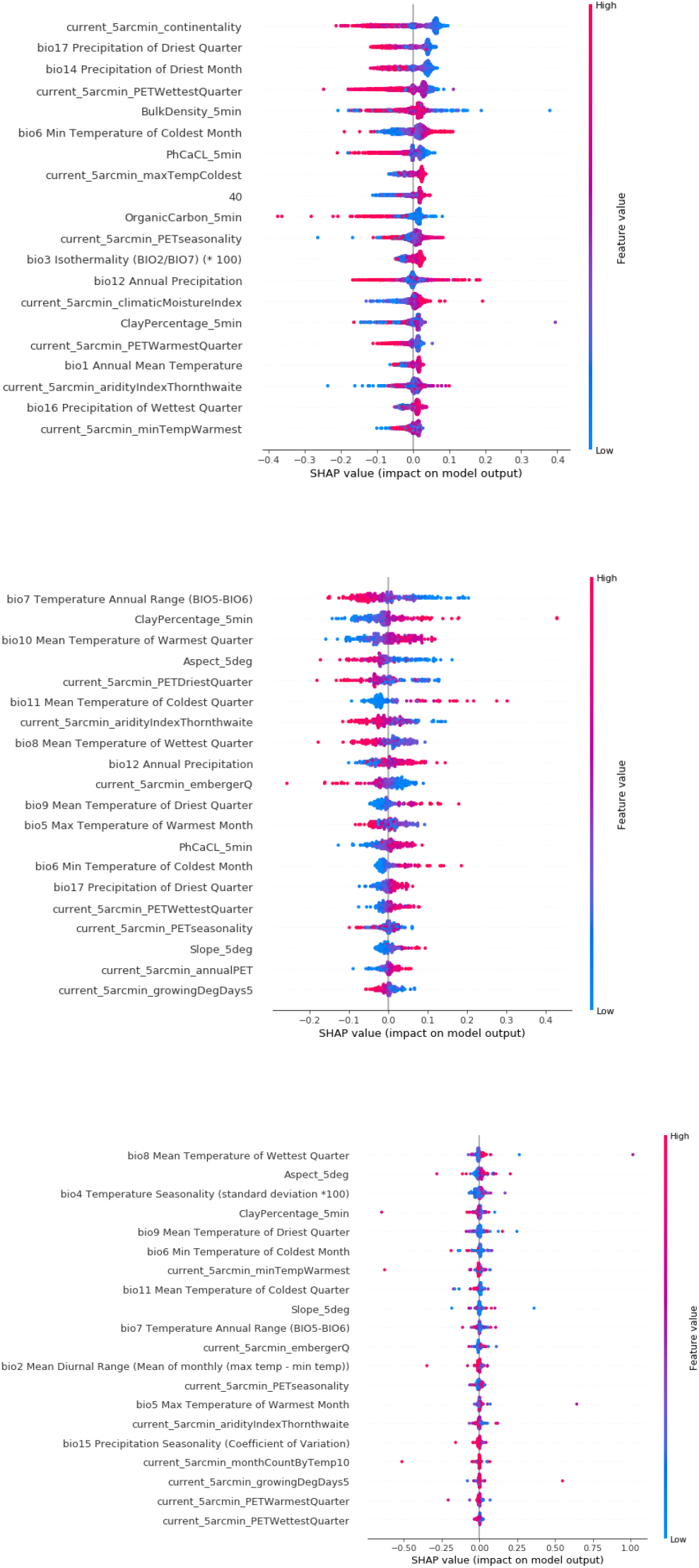

#### D Occurrence maps pilot and extended models

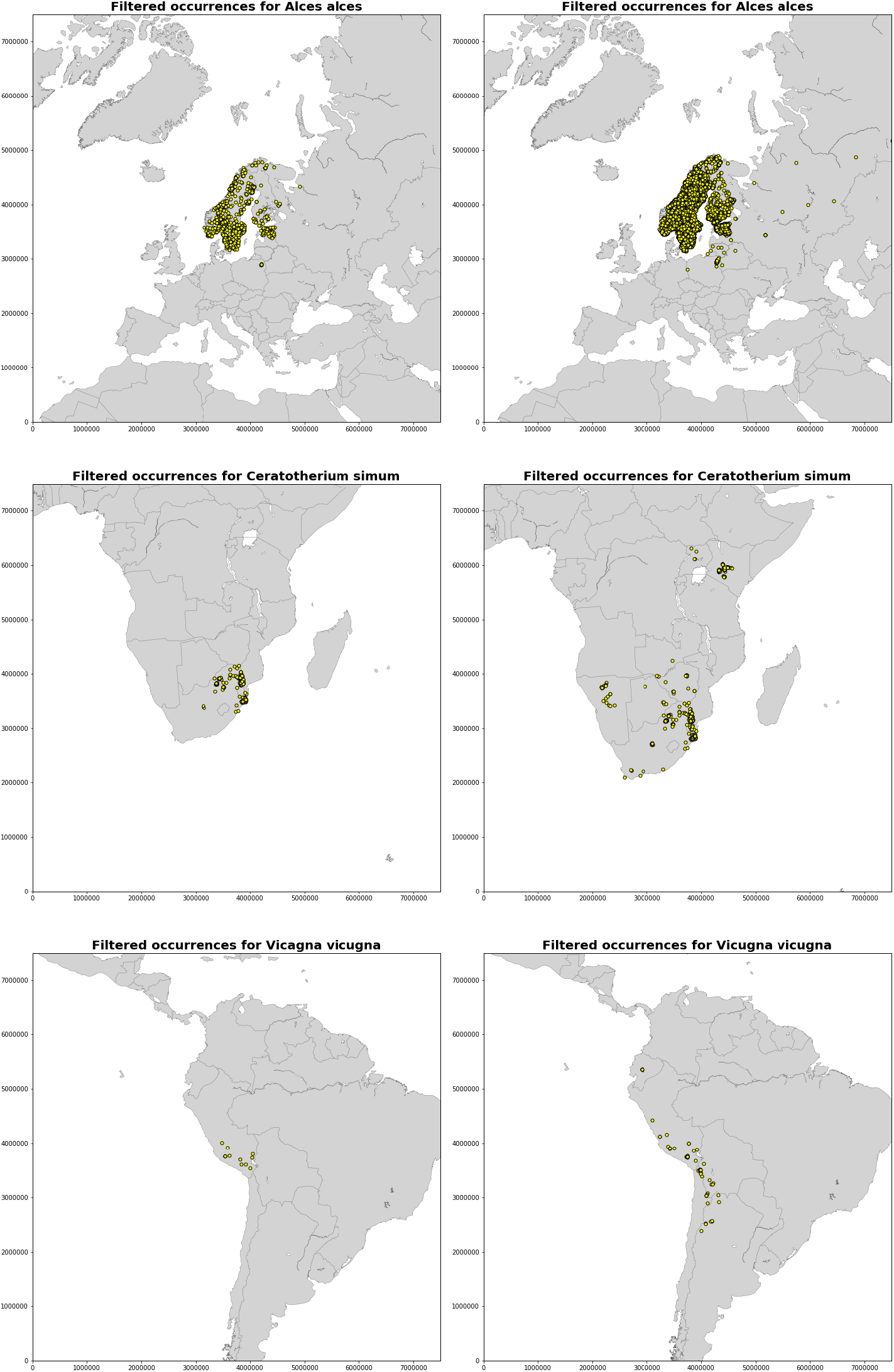

#### E Information trade-off DNN

##### 1.

Graph from Wolchover (*90*) based on research by Schwartz & Tishby (*57*), showing the internal trade-off in DNN between efficient representation of information from input features and maintaining predictive capabilities during training.

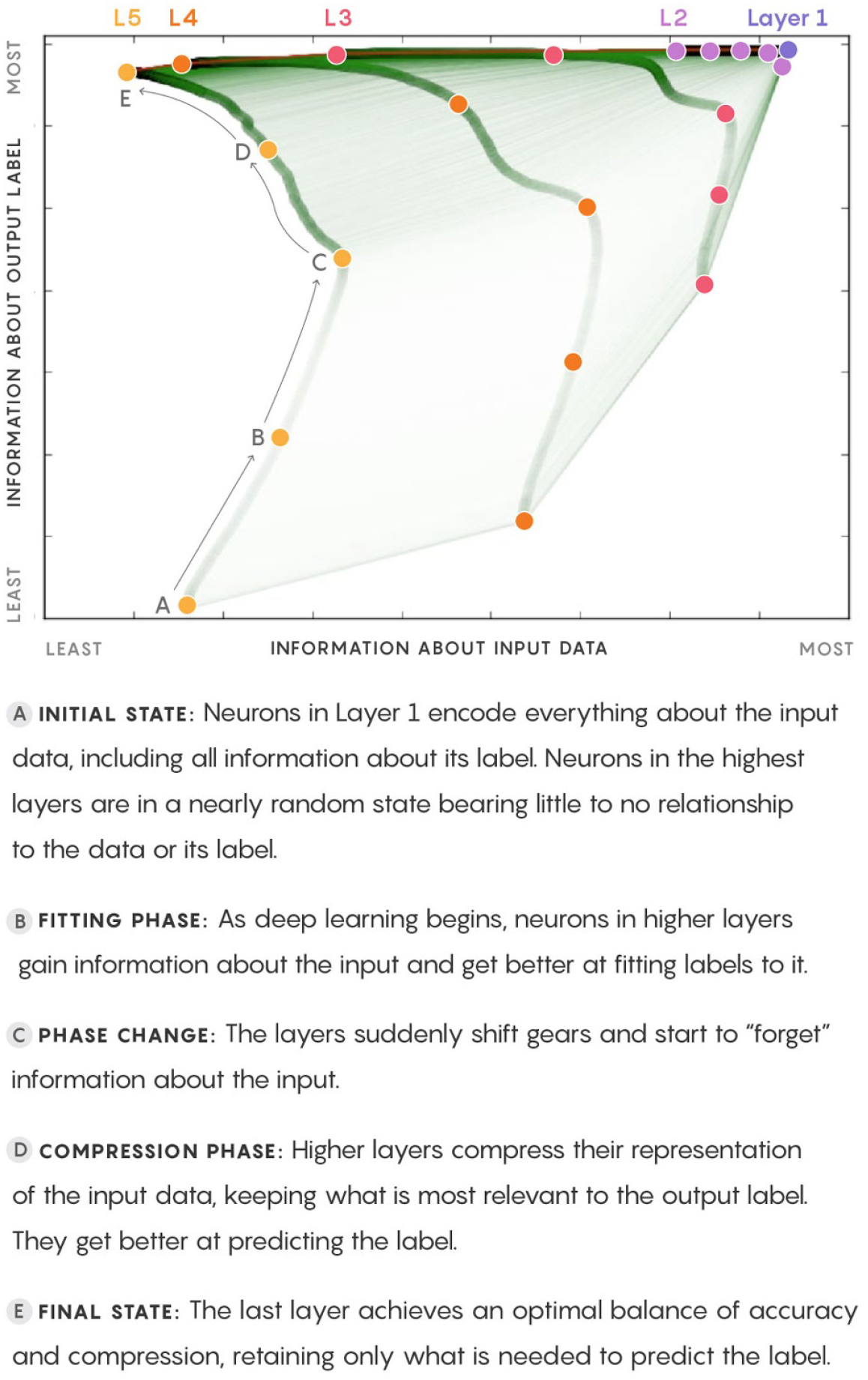

##### 2.

Graph from Saxe (*55*), with left graph representing a replication from the study of Schwartz & Tishby and right a replication using ReLu activation functions instead of tanh and sigmoid. The mutual information now increases linearly in most layers except for the final layer, and there is no clear trade-off observed.

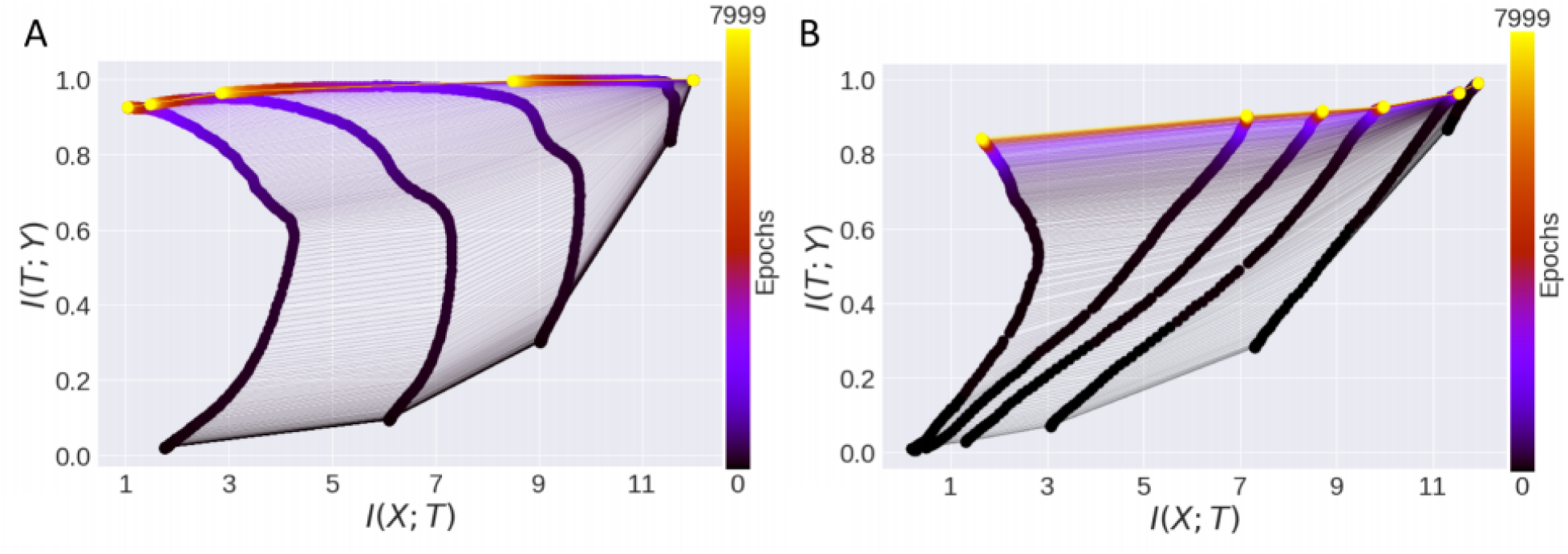

## Footnotes

1 All data and code are publicly available on the project’s online github repository: https://github.com/naturalis/trait-geo-diverse-dl

## References

(1) Christopher N Johnson et al. “Biodiversity losses and conservation responses in the Anthropocene”. In: Science 356.6335 (2017), pp. 270–275.

(2) Xingli Giam. “Global biodiversity loss from tropical deforestation”. In: Proceedings of the National Academy of Sciences 114.23 (2017), pp. 5775–5777.

(3) A Benıtez-López et al. “The impact of hunting on tropical mammal and bird populations”. In: Science 356.6334 (2017), pp. 180–183.

(4) Georgina M Mace et al. “Aiming higher to bend the curve of biodiversity loss”. In: Nature Sustainability 1.9 (2018), p. 448.

(5) Camilo Mora et al. “How many species are there on Earth and in the ocean?” In: PLoS biology 9.8 (2011), e1001127.

(6) Hans Ter Steege et al. “Hyperdominance in the Amazonian tree flora”. In: Science 342.6156 (2013), p. 1243092.

(7) Thomas J Matthews and Robert J Whittaker. “On the species abundance distribution in applied ecology and biodiversity management”. In: Journal of Applied Ecology 52.2 (2015), pp. 443–454.

(8) Mark J Costello, Robert M May, and Nigel E Stork. “Can we name Earth’s species before they go extinct?” In: science 339.6118 (2013), pp. 413–416.

(9) Robert J Whittaker et al. “Conservation biogeography: assessment and prospect”. In: Diversity and distributions 11.1 (2005), pp. 3–23.

(10) Antoine Guisan and Niklaus E Zimmermann. “Predictive habitat distribution models in ecology”. In: Ecological modelling 135.2-3 (2000), pp. 147–186.

(11) Antoine Guisan and Wilfried Thuiller. “Predicting species distribution: offering more than simple habitat models”. In: Ecology letters 8.9 (2005), pp. 993–1009.

(12) Gurutzeta Guillera-Arroita. “Modelling of species distributions, range dynamics and communities under imperfect detection: advances, challenges and opportunities”. In: Ecography 40.2 (2017), pp. 281–295.

(13) Gurutzeta Guillera-Arroita et al. “Is my species distribution model fit for purpose? Matching data and models to applications”. In: Global Ecology and Biogeography 24.3 (2015), pp. 276–292.

(14) André SJ van Proosdij et al. “Minimum required number of specimen records to develop accurate species distribution models”. In: Ecography 39.6 (2016), pp. 542–552.

(15) Steven J Phillips, Robert P Anderson, and Robert E Schapire. “Maximum entropy modeling of species geographic distributions”. In: Ecological modelling 190.3-4 (2006), pp. 231–259.

(16) Steven J Phillips and Miroslav Dudık. “Modeling of species distributions with Maxent: new extensions and a comprehensive evaluation”. In: Ecography 31.2 (2008), pp. 161–175.

(17) Cory Merow, Matthew J Smith, and John A Silander Jr. “A practical guide to MaxEnt for modeling species’ distributions: what it does, and why inputs and settings matter”. In: Ecography 36.10 (2013), pp. 1058–1069.

(18) William Fithian and Trevor Hastie. “Finite-sample equivalence in statistical models for presence-only data”. In: The annals of applied statistics 7.4 (2013), p. 1917.

(19) J Andrew Royle et al. “Likelihood analysis of species occurrence probability from presence-only data for modelling species distributions”. In: Methods in Ecology and Evolution 3.3 (2012), pp. 545–554.

(20) Jane Elith et al. “A statistical explanation of MaxEnt for ecologists”. In: Diversity and distributions 17.1 (2011), pp. 43–57.

(21) K Remya, A Ramachandran, and S Jayakumar. “Predicting the current and future suitable habitat distribution of Myristica dactyloides Gaertn. using MaxEnt model in the Eastern Ghats, India”. In: Ecological engineering 82 (2015), pp. 184–188.

(22) Magdalena Bennett et al. “Shifts in habitat suitability and the conservation status of the Endangered Andean cat Leopardus jacobita under climate change scenarios”. In: Endangered Species Research 16 (2017), pp. 283–294.

(23) Yoav Shoham et al. The AI Index 2018 Annual Report. 2018.

(24) Jürgen Schmidhuber. “Deep learning in neural networks: An overview”. In: Neural networks 61 (2015), pp. 85–117.

(25) Yanming Guo et al. “Deep learning for visual understanding: A review”. In: Neurocomputing 187 (2016), pp. 27–48.

(26) Aurélien Géron. “Hands-on machine learning with Scikit-Learn and TensorFlow: concepts, tools, and techniques to build intelligent systems”. In: O’Reilly Media, Inc., 2017. Chap. 10: Introduction to Artificial Neural Networks.

(27) Ian Goodfellow, Yoshua Bengio, and Aaron Courville. Deep learning. MIT press, 2016.

(28) Qinghua Guo and Yu Liu. “ModEco: an integrated software package for ecological niche modeling”. In: Ecography 33.4 (2010), pp. 637–642.

(29) Mauro Enrique de Souza Munoz et al. “openModeller: a generic approach to species’ potential distribution modelling”. In: GeoInformatica 15.1 (2011), pp. 111–135.

(30) Shinji Fukuda et al. “Habitat prediction and knowledge extraction for spawning European grayling (Thymallus thymallus L.) using a broad range of species distribution models”. In: Environmental modelling & software 47 (2013), pp. 1–6.

(31) Xinhai Li and Yuan Wang. “Applying various algorithms for species distribution modelling”. In: Integrative Zoology 8.2 (2013), pp. 124–135.

(32) David J Harris. “Generating realistic assemblages with a joint species distribution model”. In: Methods in Ecology and Evolution 6.4 (2015), pp. 465–473.

(33) MIT. MIT 6.S191 Introduction to Deep Learning. http://introtodeeplearning.com/materials/2019_6S191_L1.pdf. 2019

(34) Brooke L Bateman et al. “Biotic interactions influence the projected distribution of a specialist mammal under climate change”. In: Diversity and Distributions 18.9 (2012), pp. 861–872.

(35) Phillip PA Staniczenko et al. “Linking macroecology and community ecology: refining predictions of species distributions using biotic interaction networks”. In: Ecology letters 20.6 (2017), pp. 693–707.

(36) Carsten F Dormann et al. “Collinearity: a review of methods to deal with it and a simulation study evaluating their performance”. In: Ecography 36.1 (2013), pp. 27–46.

(37) Niels Raes and Jesús Aguirre-Gutiérrez. “Modeling Framework to Estimate and Project Species Distributions Space and Time”. In: Mountains, Climate and Biodiversity (2018), p. 309.

(38) Thomas Kluyver et al. “Jupyter Notebooks-a publishing format for reproducible computational workflows.” In: ELPUB. 2016, pp. 87–90.

(39) Allen Downey et al. How To Think Like A Computer Scientist: Learning with Python 3. https://buildmedia.readthedocs.org/media/pdf/howtothink/latest/howtothink.pdfs. 2016.

(40) Elke Hendrix and Rutger Vos. “Differentiation between wild and domesticated Ungulates based on ecological niches”. In: bioRxiv Preprint (2019).

(41) Donald Knuth. The Global Biodiversity Information Facility (2019): What is GBIF? url: https://www.gbif.org/what-is-gbif. (accessed: 03.04.2019).

(42) Stephen E Fick and Robert J Hijmans. “WorldClim 2: new 1-km spatial resolution climate surfaces for global land areas”. In: International journal of climatology 37.12 (2017), pp. 4302–4315.

(43) Pascal O Title and Jordan B Bemmels. “ENVIREM: an expanded set of bioclimatic and topographic variables increases flexibility and improves performance of ecological niche modeling”. In: Ecography 41.2 (2018), pp. 291–307.

(44) Wei Shangguan et al. “A global soil data set for earth system modeling”. In: Journal of Advances in Modeling Earth Systems 6.1 (2014), pp. 249–263.

(45) Morgane Barbet-Massin et al. “Selecting pseudo-absences for species distribution models: how, where and how many?” In: Methods in ecology and evolution 3.2 (2012), pp. 327–338.

(46) IUCN. IUCN Red List of Threathened Species (2019): Spatial Data: Terrestrial Mammals. url: https://www.iucnredlist.org/resources/spatial-data-download. (accessed: 31.05.2019).

(47) David M Olson et al. “Terrestrial Ecoregions of the World: A New Map of Life on Earth. A new global map of terrestrial ecoregions provides an innovative tool for conserving biodiversity”. In: BioScience 51.11 (2001), pp. 933–938.

(48) Jonathan M Hoekstra et al. The Atlas of Global Conservation. Vol. 67. University of California Press Berkeley, CA, 2010.

(49) Francois Chollet et al. Keras. https://github.com/fchollet/keras. 2015.

(50) Guillaume Lemaıtre, Fernando Nogueira, and Christos K Aridas. “Imbalanced-learn: A python toolbox to tackle the curse of imbalanced datasets in machine learning”. In: The Journal of Machine Learning Research 18.1 (2017), pp. 559–563.

(51) Scott Lundberg. SHAP. https://shap.readthedocs.io/en/latest/#. 2018.

(52) Lloyd S Shapley. “A value for n-person games”. In: Contributions to the Theory of Games 2.28 (1953), pp. 307–317.

(53) Scott M Lundberg and Su-In Lee. “A unified approach to interpreting model predictions”. In: Advances in Neural Information Processing Systems. 2017, pp. 4765–4774.

(54) Avanti Shrikumar, Peyton Greenside, and Anshul Kundaje. “Learning important features through propagating activation differences”. In: Proceedings of the 34th International Conference on Machine Learning-Volume 70. JMLR. org. 2017, pp. 3145–3153.

(55) Andrew Michael Saxe et al. “On the information bottleneck theory of deep learning”. In: (2018).

(56) Naftali Tishby and Noga Zaslavsky. “Deep learning and the information bottleneck principle”. In: 2015 IEEE Information Theory Workshop (ITW). IEEE. 2015, pp. 1–5.

(57) Ravid Schwartz-Ziv and Naftali Tishby. “Opening the black box of deep neural networks via information”. In: arXiv preprint arXiv:1703.00810 (2017).

(58) Guanhua Zheng, Jitao Sang, and Changsheng Xu. “Understanding deep learning generalization by maximum entropy”. In: arXiv preprint arXiv:1711.07758 (2017).

(59) Sarunas J Raudys and Anil K. Jain. “Small sample size effects in statistical pattern recognition: Recommendations for practitioners”. In: IEEE Transactions on Pattern Analysis & Machine Intelligence 3 (1991), pp. 252–264.

(60) Sinno Jialin Pan and Qiang Yang. “A survey on transfer learning”. In: IEEE Transactions on knowledge and data engineering 22.10 (2009), pp. 1345–1359.

(61) Deepak Soekhoe, Peter Van Der Putten, and Aske Plaat. “On the impact of data set size in transfer learning using deep neural networks”. In: International Symposium on Intelligent Data Analysis. Springer. 2016, pp. 50–60.

(62) Ross Girshick et al. “Rich feature hierarchies for accurate object detection and semantic segmentation”. In: Proceedings of the IEEE conference on computer vision and pattern recognition. 2014, pp. 580–587.

(63) Maxime Oquab et al. “Learning and transferring mid-level image representations using convolutional neural networks”. In: Proceedings of the IEEE conference on computer vision and pattern recognition. 2014, pp. 1717–1724.

(64) Jason Yosinski et al. “How transferable are features in deep neural networks?” In: Advances in neural information processing systems. 2014, pp. 3320–3328.

(65) Chen Huang et al. “Learning Deep Representation for Imbalanced Classification”. In: The IEEE Conference on Computer Vision and Pattern Recognition (CVPR). June 2016.

(66) Aili Qin et al. “Maxent modeling for predicting impacts of climate change on the potential distribution of Thuja sutchuenensis Franch., an extremely endangered conifer from southwestern China”. In: Global Ecology and Conservation 10 (2017), pp. 139–146.

(67) Keliang Zhang et al. “Maxent modeling for predicting the potential geographical distribution of two peony species under climate change”. In: Science of the Total Environment 634 (2018), pp. 1326–1334.

(68) Marcin K Dyderski et al. “How much does climate change threaten European forest tree species distributions?” In: Global change biology 24.3 (2018), pp. 1150–1163.

(69) Rafal Jozefowicz et al. “Exploring the limits of language modeling”. In: arXiv preprint arXiv:1602.02410 (2016).

(70) Tomáš Mikolov et al. “Recurrent neural network based language model”. In: Eleventh annual conference of the international speech communication association. 2010.

(71) Stephan Rasp, Michael S Pritchard, and Pierre Gentine. “Deep learning to represent subgrid processes in climate models”. In: Proceedings of the National Academy of Sciences 115.39 (2018), pp. 9684–9689.

(72) Sangmok Lee and Donghyun Lee. “Improved prediction of harmful algal blooms in four Major South Korea’s Rivers using deep learning models”. In: International journal of environmental research and public health 15.7 (2018), p. 1322.

(73) Maarten Schermer and Laurens Hogeweg. “Supporting Citizen Scientists with Automatic Species Identification using Deep Learning Image Recognition Models”. In: Joint meeting of the Society for the Preservation of Natural History Collections (SPNHC) and Biodiversity Information Standards (TDWG), Dunedin, New Zealand. 2018. url: https://drive.google.com/file/d/1xAB8NmwVlqwNivzcXlcprJXvnWpc0Hhv/view.

(74) Maarten Schermer, Laurens Hogeweg, and Max Caspers. “Using Deep Learning in Collection Management to Reduce the Taxonomist’s Workload”. In: Joint meeting of the Society for the Preservation of Natural History Collections (SPNHC) and Biodiversity Information Standards (TDWG), Dunedin, New Zealand. 2018. url: https://drive.google.com/file/d/1JBAg0GVcLRElvtWQ8H5wXvVWltWSIvhg/view.

(75) Tom Fawcett. “An introduction to ROC analysis”. In: Pattern recognition letters 27.8 (2006), pp. 861–874.

(76) Seong Ho Park, Jin Mo Goo, and Chan-Hee Jo. “Receiver operating characteristic (ROC) curve: practical review for radiologists”. In: Korean Journal of Radiology 5.1 (2004), pp. 11–18.

(77) D Richard Baughman and Yih An Liu. Neural networks in bioprocessing and chemical engineering. Academic press, 1995.

(78) Hao Li et al. “Visualizing the loss landscape of neural nets”. In: Advances in Neural Information Processing Systems. 2018, pp. 6389–6399.

(79) Sagar Sharma. Activation Functions in Neural Networks. https://towardsdatascience.com/activation-functions-neural-networks-1cbd9f8d91d6. 2015.

(80) Arunava. Derivative of the Sigmoid function. https://towardsdatascience.com/derivative-of-the-sigmoid-function-536880cf918e. 2018.

(81) Alex Smola and SVN Vishwanathan. “Introduction to machine learning”. In: Cambridge University, UK 32 (2008), p. 34.

(82) Boris T Polyak. “Some methods of speeding up the convergence of iteration methods”. In: USSR Computational Mathematics and Mathematical Physics 4.5 (1964), pp. 1–17.

(83) Ilya Sutskever et al. “On the importance of initialization and momentum in deep learning”. In: International conference on machine learning. 2013, pp. 1139–1147.

(84) John Duchi, Elad Hazan, and Yoram Singer. “Adaptive subgradient methods for online learning and stochastic optimization”. In: Journal of Machine Learning Research 12.Jul (2011), pp. 2121–2159.

(85) Sebastian Ruder. “An overview of gradient descent optimization algorithms”. In: arXiv preprint arXiv:1609.04747 (2016).

(86) Diederik P Kingma and Jimmy Ba. “Adam: A method for stochastic optimization”. In. arXiv preprint arXiv:1412.6980 (2014).

(87) Gareth James et al. An introduction to statistical learning. Vol. 112. Springer, 2013.

(88) Nitish Srivastava et al. “Dropout: a simple way to prevent neural networks from overfitting”. In: The Journal of Machine Learning Research 15.1 (2014), pp. 1929–1958.

(89) Cody Marie Wild. One Feature Attribution Method to (Supposedly) Rule Them All: Shapley Values. https://towardsdatascience.com/one-feature-attribution-method-to-supposedly-rule-them-all-shapley-values-f3e04534983d. 2018.

(90) Natalie Wolchover and Lucy Reading. “New theory cracks open the black box of deep learning”. In: Quanta Magazine 3 (2017). url: https://www.quantamagazine.org/new-theory-cracks-open-the-black-box-of-deep-learning-20170921/.

